# Deep mutational scanning reveals the functional constraints and evolutionary potential of the influenza A virus PB1 protein

**DOI:** 10.1101/2023.08.27.554986

**Authors:** Yuan Li, Sarah Arcos, Kimberly R. Sabsay, Aartjan J.W. te Velthuis, Adam S. Lauring

## Abstract

The influenza virus polymerase is central to influenza virus evolution. Adaptive mutations within the polymerase are often a prerequisite for efficient spread of novel animal-derived viruses in human populations. The polymerase also determines fidelity, and therefore the rate at which the virus will acquire mutations that lead to host range expansion, drug resistance, or antigenic drift. Despite its importance to viral replication and evolution, our understanding of the mutational effects and associated constraints on the influenza RNA-dependent RNA polymerase (RdRp) is relatively limited. We performed deep mutational scanning of the A/WSN/1933(H1N1) PB1, generating a library of 95.4% of amino acid substitutions at 757 sites. After accuracy filters, we were able to measure replicative fitness for 13,354 (84%) of all possible amino acid substitutions, and 16 were validated by results from pairwise competition assays. Functional and structural constraints were better revealed by individual sites involved in RNA or protein interactions than by major subdomains defined by sequence conservation. Mutational tolerance, as defined by site entropy, was correlated with evolutionary potential, as captured by diversity in available H1N1 sequences. Of 29 beneficial sites, many have either been identified in the natural evolution of PB1 or shown experimentally to have important impacts on replication and adaptation. Accessibility of amino acid substitutions by single nucleotide mutation was a key factor in determining whether mutations appeared in natural PB1 evolution. Our work provides a comprehensive map of mutational effects on a viral RdRp and a valuable resource for subsequent studies of influenza replication and evolution.

## Introduction

Viral RNA-dependent RNA polymerases (RdRp) are central to RNA virus replication and evolution. The RdRp replicates the genome and is a key determinant for replicative fitness and viral mutation rates. For negative-strand RNA viruses, the RdRp is also responsible for transcription, thereby regulating protein expression. Mutations within the RdRp influence host adaptation (1, 2, 3), replication fidelity (4), post-translational modifications (5), and host immune responses (6, 7).

The evolution of viral RdRp is functionally and structurally constrained. Functional constraints include requirements for interactions with RNAs and other proteins, adaptation to new replication environments (8), the deleterious impact of low fidelity (4), and viral codon abundance (9, 10, 11). Residues that are involved in obligatory interactions tend to be less tolerant to mutation and evolve at a slower rate (12-15). The primary structural constraints are solvent accessibility (16), maintenance of molecular flexibility (17, 18, 19), intermolecular interactions (20, 21), and key protein secondary structures (22). For example, the establishment of secondary structures requires certain biochemical characteristics conferred by a limited number of amino acids (22), and mutations in buried residues often have a bigger fitness effect, as their change will impact nearby residues (23, 24).

The Influenza virus RdRp is a heterotrimer that consists of three subunits: polymerase basic 1 (PB1), polymerase basic 2 (PB2), and polymerase acidic (PA), in which PB1 functions as the catalytic subunit. The PB1 subunit may have additional functional and structural constraints, because it cooperates with the two other polymerase subunits and viral nucleoproteins (NP) in transcription and genome replication. During transcription, PB1 guides the capped primer cleaved from a host pre-mRNA by PB2 and PA into the polymerase active site and stabilizes it on the 3′ end of the viral RNA (vRNA) template in the active site (25). The PB1 RdRp then extends the capped primer through the incorporation of nucleoside triphosphates, separates the template-product duplex downstream of the active site, and extrudes the viral mRNA through the product exit channel and the copied template through the template exit channel (25). The interactions among PB1, PB2, and PA shift at every stage of transcription (26). During replication, a vRNA is copied into a complementary RNA (cRNA). Next, the cRNA product serves as the template for negative strand vRNA synthesis. The process of vRNA and cRNA synthesis not only requires the coordination of polymerase subunits, but also interactions with an encapsidating RdRp and host protein ANP32 to form an RdRp dimer, a trans-activating RdRp to induce correct replication initiation, conformational changes to transfer the nascent vRNA or cRNA to the additional RdRp, and recruitment of viral nucleoprotein to encapsidate the nascent vRNA and cRNA molecules (27).

Given the importance of PB1 to influenza virus replication and evolution, defining the fitness effects of amino acid substitutions can elucidate the relevant functional and structural constraints. Deep mutational scanning (DMS) – saturation mutagenesis combined with deep sequencing – is a massively parallel approach that has recently been used to explore the fitness landscapes of viral proteins (15, 28-31). Here, we applied deep mutational scanning to the influenza virus A/WSN/1933(H1N1) (abbreviated WSN33) PB1 RdRp subunit, identifying constrained regions of the protein and relating beneficial mutations to those observed in natural evolution. Overall, our study provides a comprehensive resource for studies of influenza virus replication and evolution.

## Methods

### Cell lines and media

MDCK-SIAT1-TMPRSS2 and HEK293T-CMV-PB1 were provided by Dr. Jesse Bloom (Fred Hutchinson Cancer Research Center) and maintained in D10 media (Dulbecco’s Modified Eagle Medium, DMEM, Invitrogen, 11995-065), with 10% heat inactivated Fetal Bovine Serum (FBS, Gibco, 26140-079), 1% L-Glutamine (100×, Gibco, 25030-081), and 1% Pen+Strep (10,000 U/mL P, 10,000 μg/mL S, Invitrogen, 15140-122)). A549 cells were maintained in A549 growth media (DMEM, high glucose, with L-glutamine, without Na pyruvate (Invitrogen 11965-092), with 10% FBS, 1% Pen+Strep, 0.1875% Bovine albumin fraction V (7.5%, Invitrogen 15260-037), and 2.5% HEPES (1M, Invitrogen, 15630-080). We used IGM+ media (Opti-MEM1 Reduced Serum Media (Gibco, 31985-070), with 0.5% heat inactivated FBS, 1% Pen+Strep, 0.3% Bovine albumin fraction V, and 500 μL of 100mg/ml CaCl_2_) for 24 hours following transfection. We used WNM media (Medium 199, Gibco, 11043-023, no phenol red), 0.5% heat inactivated FBS, 1% Pen+Strep, 0.3% Bovine albumin fraction V, 2.5% HEPES, and 500 μL of 100mg/ml CaCl_2_) for TCID_50_ assays. We used A549 growth media for seeding cells for viral passages and A549 viral media (DMEM, high glucose, with L-glutamine, without Na pyruvate (Invitrogen 11965-092), with 1% Pen+Strep, 0.1875% Bovine albumin fraction V, 2.5% HEPES, and TPCK-trypsin at a final concentration of 4 μg/ml) for virus infections.

### Construction of PB1 codon mutant plasmid libraries

PB1 codon mutant libraries were generated using an overlapping PCR strategy described in (28) with (32) as a reference. We used code in (33) first described by (34) with the modifications from (35) to generate tiled primers for mutagenesis and code from (36) to determine how library diversity would be impacted by restriction enzymes used in cloning. We performed 10 cycles of fragment PCR (Round one) with 1.2 µg of plasmid (pHW2000) containing the wildtype PB1 sequence from WSN33, and 20 cycles of joining PCR (Round two). The lengths of PCR products were checked by gel electrophoresis. In a pilot experiment in which we generated PB1 variants for 96 out of the 758 sites, we randomly picked PCR products from 24 clones for Sanger sequencing to evaluate the library mutation rate. Twenty out of 24 clones had only a single codon mutation at the target site, and 4 clones were wild type.

We pooled an equal volume from the 758 PCR reactions into 16 pools. Each pool was digested by restriction enzyme *Aar*I and the 16 pools combined into one variant insert pool. We used T4 DNA ligase (NEB, #M0202L) to ligate the variant insert pool into BsmBI-digested pHW2000 plasmid and transformed Stellar™ Competent Cells (Takara, #636763) according to the manufacturer’s instructions. We independently performed the ligation and transformation three times to create three libraries. We plated the transformed cells onto Nunc™ Square BioAssay Dishes (Thermo Scientific, #240845) and obtained 82,800 - 118,800 colonies for each library replicate. Plasmid DNA was extracted directly from the pooled colonies using a QIAGEN Plasmid Maxi Kit (QIAGEN, #12162).

### Transfection

We generated variant virus libraries by transfecting HEK293T-CMV-PB1 cells, which constitutively express the wild type PB1 protein from WSN33. For each variant plasmid library, we seeded 36 wells of 6-well plates with 5×10^5^ MDCK-SIAT1-TMPRSS2 cells and 5×10^5^ HEK293T-CMV-PB1 cells. Seventeen hours later, we transfected each well with 1 μg each of the seven plasmids containing the seven wildtype WSN33 genome segments and the 1 µg of the PB1 variant library using TransIT®-LT1 Transfection Reagent (MIR 2300). We used the same procedure to make the wild type WSN33 viruses as a control, only on a smaller scale (6 wells) and using the wildtype WSN33 PB1 in place of the variant plasmid library. At 24 hours post transfection, we replaced the transfection media with fresh IGM+, then incubated for an additional 24 hours. At 48 hours post transfection, we harvested viral supernatants by centrifuging at 200 ×g for 5 minutes. Three virus variant libraries and the wild type virus control were aliquoted and snap-frozen in 0.5% glycerol prior to storage at - 80℃.

### Determination of virus titer

Viruses were titered by median Tissue Culture Infective Dose (TCID_50_) on MDCK-SIAT1-TMPRSS2 cells. For each assay, we seeded 6×10^3^ MDCK-SIAT1-TMPRSS2 cells in 100 μL of WNM media in each well of a 96-well plate. Seventeen hours later, we serially diluted the virus samples 1:10 with WNM media supplemented with 4 μg/mL TPCK-trypsin reconstituted in PBS to 1 mg/mL for working stock and added 100 μL virus per well. We incubated the plates at 37℃ and monitored daily for CPE up to four days.

### Viral passages

Each passage had 1×10^6^ infectious viral particles on 1×10^8^ A549 cells to achieve an approximate multiplicity of infection (MOI) of 0.01 TCID_50_/cell. We seeded 8×10^7^ A549 cells in a total of 60 mL A549 growth media in three T182 flasks. Seventeen hours later, we suspended 1×10^6^ TCID of virus in 45 mL of A549 viral media with 4 μg/mL freshly added TPCK-trypsin. We aspirated the overnight A549 growth media, rinsed the cells gently with pre-warmed PBS, and added 15 mL of viral dilution to each flask. Three hours after infection, we removed the inoculum, rinsed the cells again with pre-warmed PBS, and replaced the inoculum with 20 mL of fresh A549 viral media per flask with 4 μg/mL TPCK-trypsin. We harvested viral supernatants by centrifugation at 400 ×g for 4 minutes 48 hours after infection and snap-froze the supernatant in 0.5% glycerol prior to storage at -80℃.

### Barcoded subamplicon sequencing

Passaged viruses were concentrated by ultracentrifugation at 27,000 rpm, using Thermo Scientific Sorvall WX Ultra Series Centrifuge with rotor Sorvall AH-629 (DuPont Instruments), for 2 hours at 4℃ using Beckman Coulter Centrifuge Tubes (25×89 mm, 344058). We then resuspended the viruses in 500 μL of residual media and extracted viral RNA using a QIAamp Viral RNA Mini Kit (Qiagen, 52906). To accurately measure mutation frequencies, we used a barcoded-subamplicon sequencing strategy described in (37) that adds unique sequence barcodes to every DNA molecule in a sample, as follows.

We reverse transcribed the extracted RNA using SuperScript™ III First-Strand Synthesis System (Invitrogen, 18080-051) and performed PCR to amplify the entire PB1 open reading frame (PCR0). For plasmid samples, we used 2 μL of plasmid DNA at 10 ng/μL as template in PCR0. We cleaned up the PCR0 products using GeneJet PCR clean up kit (GeneJet, K0702) and gel-isolated the bands corresponding to full PB1 genome length (∼2341 bp).

Next, we PCR amplified the PB1 gene in eight sub-amplicons (PCR1). The subamplicons were designed to start and end in full codons, and each subamplicon starts precisely after the previous subamplicon ends. In this way, the nucleotides in one codon in a PB1 DNA molecule will only be calculated once. Forward and reverse primers for PCR1 contained random 8N barcodes at their 5⍰ termini to uniquely label every cDNA molecule in the template. Theoretically, there would be 4^16^ = 4.29 x 10^9^ unique barcodes. The template input for PCR1 was limited to ∼8 x 10^7^ molecules such that each was uniquely barcoded. Illumina compatible, sample-specific adapters were added in a subsequent PCR reaction, PCR2. Eight subamplicons for each sample were pooled together, and we used ∼1x 10^6^ uniquely barcoded molecules from PCR1 as template and unique dual (UD) indexed primers to diminish the issue of index hopping. Finally, we gel-isolated the PCR2 products before sequencing on an Illumina NextSeq 1000, P2 600 cycle (2 x 300 PE), with 20% PhiX. We conducted two sequencing runs with 60 μL of the combined PCR2 products at 5 nM, 30 μL for each run, and merged the reads for analysis. We used KOD Hot Start Master Mix (EMD Millipore, 71842) to perform all PCRs. Primers and cycling programs can be found in a Supplementary File (Supplemental Text).

### Analysis of deep sequencing data

Sequence files were analyzed using dms_tools2 (38), which groups the paired-end reads with the same PCR1 barcodes. Sequences were discarded if the Q-score of any nucleotide in the barcode was < 15. Consensus sequences were generated for barcodes with at least two reads and aligned to the reference genome to record the codon at each site for that molecule.

We calculated the fitness of each mutation based on the enrichment ratio method described in (31) with modifications. We calculated the frequency of mutation *i* at site *s* as:

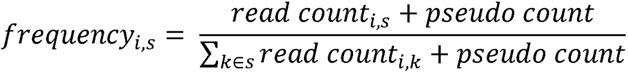

where the pseudo count was added to ensure a non-zero denominator and fixed as 1 by default. To off-set frequency inflation by the pseudo count, we discarded a mutation if its read count in the variant plasmid library was less than 10. We then discarded the mutations whose frequency in the variant plasmid library was not at least 6-fold higher than that in the wildtype plasmid library. With these filters, we calculated the enrichment ratio as:

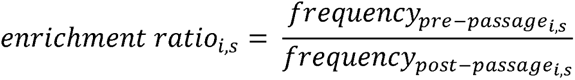

and we defined “fitness” as log_1O_ (*enrichment ratio*) normalized by the average fitness of silent mutations in corresponding amplicons:

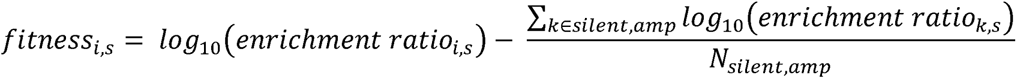

The fitness of a mutation at a certain site used for subsequent analyses was the average fitness of that in all replicates where it was available.

### Analysis of naturally occurring influenza sequences

We downloaded the influenza sequences from Global Initiative on Sharing All Influenza Data (GISAID) from 1918 to 2023, with the filtering conditions of “type A”, “H1N1”, “human host”, “required segment PB1”, and “complete sequences only”. According to CDC’s timeline for 2009 H1N1 pandemic (39), we classified pre-09 strains as all sequences collected before April 14, 2009, and post-09 strains as sequences collected after August 12, 2010. We discarded the sequences collected during the pandemic to avoid the time period when pre- and post-09 strains might be co-circulating. We downloaded the amino acid sequences along with the corresponding metadata and filtered out any sequences that had been passaged in eggs. We aligned the sequences to the wildtype WSN33 amino acid sequence using MAFFT (40). The entropy of a site was measured as the Shannon Entropy (41) of all amino acids that appeared at that site:

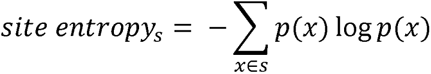

To adjust for uneven sampling over time, we adopted the weighted entropy method described in (42). Briefly, we grouped the sequences by collection year, calculated the frequencies of the amino acids in each year, and used the average of amino acid frequencies over all years for the entropy calculation.

### Protein structure visualization and analysis

We used UCSF ChimeraX (43) for protein visualizations, including movies. To visualize site entropy on the PB1 protein, we replaced the b-factor column with site entropy data in the PDB files. We identified protein-RNA contacts using LigPlot Plus with default thresholds for the maximum distance between interacting atoms (44). The protein structures used are as follows: 5D9A (apoenzyme), 7NHX (template binding, early), 6T0N (template binding, late), 5M3H (cap-snatching), 6RR7 (pre-initiation), 6QCW (mixed pre-initiation), 6QCV (mixed pre-catalysis), 6QCX (mixed post-incorporation), 6SZV (elongation), and 6SZU (termination).

### Measurement of accessible surface area

We measured the Accessible Surface Area (ASA) using PDBePISA (’Protein interfaces, surfaces and assemblies’ service PISA at the European Bioinformatics Institute. http://www.ebi.ac.uk/pdbe/prot_int/pistart.html, 45) with influenza A/Brevig Mission/1/1918(H1N1) polymerase heterotrimer structure (PDB: 7NHX). We chose to perform this and subsequent analyses with 7NHX because this is the only resolved structure for H1N1 polymerase complex, which may be a closer approximation to WSN33 RdRp. We used a default water probe of 1.4 Å in diameter to roll over the surface of the entire polymerase complex and added up all points in contact with the probe.

### Molecular dynamics simulation and measurement of root mean square fluctuation

We performed a molecular dynamics simulation of A/Brevig Mission/1/1918(H1N1) RdRp (PDB: 7NHX) to measure the relative structural flexibility of the heterotrimer. We removed the RNA molecules in the structure and modeled the missing residues (Chain B, PB1: 187-204 and 645-653) using SWISS-MODEL template based homology using full sequences from UNIPROT (PA: Q3HM39, PB1: Q3HM40, and PB2: Q3HM41). The global model quality estimate (GMQE) for this homology model is 0.88. Molecular dynamics were simulated using GROMACS on the Princeton University HPC Tiger GPU. The system build parameters used a cubic tip3p water box, charmm27 force field, neutralizing NaCl ions, temperature of 310.15 K (37℃), and time steps of 0.002 ps. The total system had 2148 protein residues, 166370 water residues and 1002 ion residues. Energy minimization was performed for a total of 100 ps and converged to a maximum force of less than 1000 kJ/mol in 2102 steps. Equilibration (both NVT and NPT ensembles) were performed for 200 ps. A 20 ns (10,000,000 steps) production simulation took roughly 19 hours. We analyzed the resulting trajectory for the root mean square fluctuations (RMSF) of the atomic positions at every time point and calculated the RMSF of each residue within the structure was from an average of RMSF values of each atom within the residue.

### Data availability

Raw sequence reads are available in the NCBI Sequence Read Archive under Bioproject #PRJNA1009589

## Results

### A comprehensive library of single amino acid substitutions in PB1

We used overlap PCR mutagenesis to create a PB1 plasmid library in which every codon in the WSN33 PB1 open reading frame is mutated to code for every other amino acid. We cloned the mutagenized plasmid library three times, independently, to make three replicate plasmid libraries (Figure 1). High depth-of-coverage sequencing demonstrated that the three plasmid libraries covered 82-93% of 24,224 possible codon mutations and 89-96% of 15,897 possible amino acid substitutions at 757 residues across 758 (stop codon included) sites in PB1 (Table 1). After excluding mutations whose frequencies might have been inflated by mutational hotspots during sequencing library preparation, each replicate library covered 64-70% of all possible amino acid substitution with 84% of all possible amino acid substitutions present in at least one replicate library.

**Figure 1.**
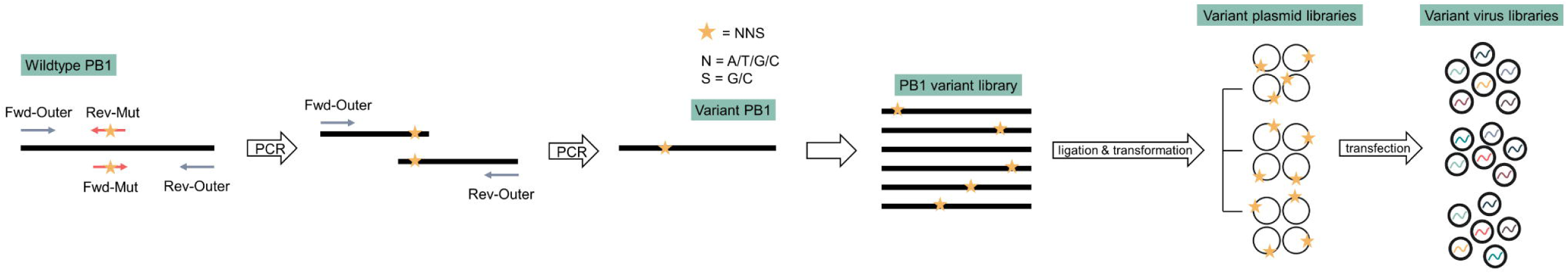
Deep mutational scanning of influenza PB1 protein. Scheme of major steps for generating variant virus libraries. We mutagenized wildtype PB1 by overlap PCR, using primers encoding *NNS* in the codon for the targeted residue. *N* refers to an equal mixture of A, T, G, and C nucleotides, while *S* refers to a mixture of only G and C. This coding is able to generate 32 codons, 20 amino acids, and stop codons. The PB1 variant library was ligated and transformed independently three times to make variant plasmid library replicates. Each plasmid library was then transfected independently along with plasmids expressing the other seven influenza segments to make three variant virus library replicates.

**Table 1.** Codon and amino acid variant diversity in plasmid libraries before and after filtering out mutations with low codon counts or under the influence of PCR errors.

|  | Library | Number of<br>codons | Percentage of<br>codons | Number of<br>amino acids | Percentage of<br>amino acids |
| --- | --- | --- | --- | --- | --- |
| Before<br>filtering | Replicate 1 | 22820 | 92.6% | 15168 | 95.4% |
|  | Replicate 2 | 21995 | 89.2% | 14786 | 93.0% |
|  | Replicate 3 | 20264 | 82.2% | 14154 | 89.0% |
| After<br>filtering | Replicate 1 | - | - | 11008 | 69.3% |
|  | Replicate 2 | - | - | 10565 | 66.5% |
|  | Replicate 3 | - | - | 10194 | 64.1% |
|  | Present in all replicates | - | - | 7351 | 46.2% |
|  | Present in at least one<br>replicate | - | - | 13354 | 84.0% |

We rescued the corresponding viral variant libraries by transfecting HEK293T cells that stably express PB1 with the plasmid libraries and bidirectional expression plasmids containing the other seven genomic segments from WSN33. The passage 0 (P0) viral stocks exhibited titers of 3.51×10^6^ to 5.27×10^7^ TCID /mL after 48 hours, slightly lower than those from “wild type” WSN33 rescues. Four hundred and eighty-two mutations in Replicate 1 (2.1% of total mutations in Replicate 1), 1371 mutations in Replicate 2 (6.2%), and 175 mutations in Replicate 3 (0.86%) were present in the plasmid library but not in the P0 viral library, which may indicate lethal mutations. The experimental lethal mutation rate was lower than the expected ∼25-30% lethal mutation fraction (46) because the wildtype PB1 protein expressed by the cells partially rescued the variant PB1 proteins with lethal mutations.

We examined the fitness effects of the mutations through serial passage of the variant virus libraries. We passaged the three libraries independently on A549 human lung epithelial carcinoma cells at an MOI of 0.01 for four passages, during which viruses carrying different PB1 substitutions competed against each other. The titers of viruses at each passage decreased slightly to 5×10^6^ to 5×10^7^ TCID /mL (Figure 2A). Forty-three percent of codons and 57% of unique amino acids on average were detected through four passages (Figure 2B). We used barcoded-subamplicon sequencing to correct for RT-PCR and sequencing errors and measured the frequencies of individual mutations in each library at passages 1 and 4 (Supplemental Figure 1). Throughout passaging, we observed signs of purifying selection, reflected by a relative reduction in the number of nonsynonymous mutations (Figure 2C) and in codons with 2 or 3 nucleotide changes (Figure 2D).

**Figure 2.**
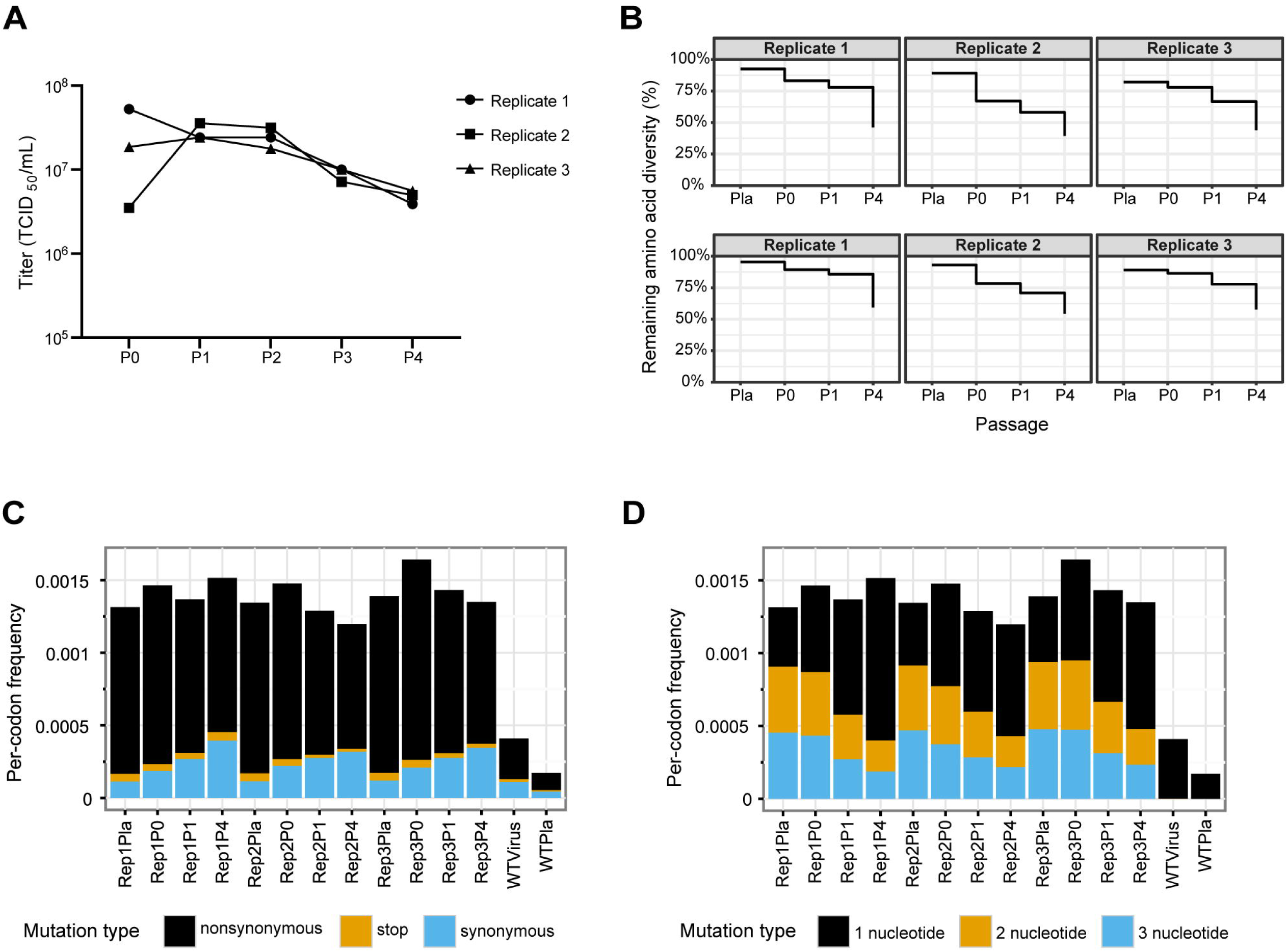
Change in codon and amino acid mutations throughout passaging. (A) Titers of variant virus libraries before and after each passage. (B) Percentage of codon and amino acid variants remaining at each passage. Pla: in plasmid library, before rescue; P0: after rescue, before passaging; P1: after the first passage; P4: after four passages. (C) Frequency of synonymous, non-synonymous, and nonsense mutations in replicate (Rep) plasmid libraries, virus libraries after passages, and the wild type (WT) plasmid and virus samples as controls. (D) Frequency of codon mutations with 1-, 2-, and 3- nucleotide change in plasmid libraries, virus libraries after passages, and the wildtype plasmid and virus samples. Frequency in both (C) and (D) panels were averaged across the PB1 gene.

### Replicative fitness of amino acid substitutions in PB1

We quantified the fitness of viral mutants at the amino acid level based on an amino acid’s frequency before and after passage (Figure 3, Supplemental Dataset). Here, the fitness of an amino acid at a site is the log_10_ enrichment ratio normalized by the average fitness of silent mutations in the same amplicon (see Methods). Since >99% of the codons at any given site in the libraries encoded the wild type amino acid, the change in frequency of wild type variants was negligible, and the measured fitness of the wild type (log_10_ of ∼1, or 0) was fixed by the experimental design. As expected, the frequency of most mutations decreased after four passages, indicating that most mutations in the influenza virus RdRp are detrimental (fitness < 0, Figure 3 and 4A). Nonsense mutations never increased in frequency. Fitness measurements were well correlated across biological replicates with Pearson correlation coefficients between 0.788 to 0.864 (Figure 4B).

**Figure 3.**
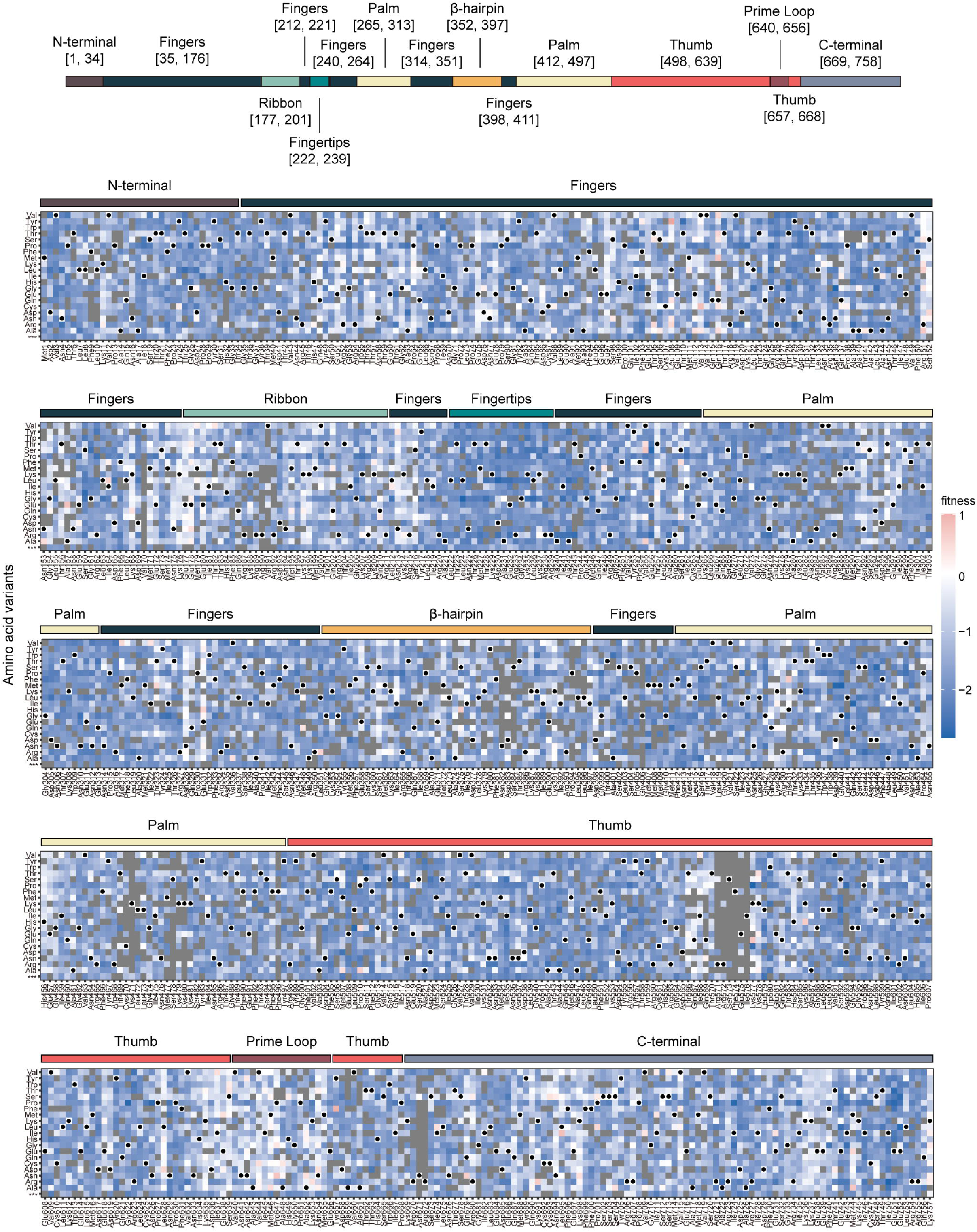
Replicative fitness of amino acid substitutions on PB1. The replicative fitness of individual amino acid variants in PB1, with subdomains annotated by the colored bar above the heatmap. Mutations in gray were excluded from the analysis due to low counts in the plasmid library or high occurrence in the wild type sample, as described in Methods. Wild type amino acids are marked by black dots.

**Figure 4.**
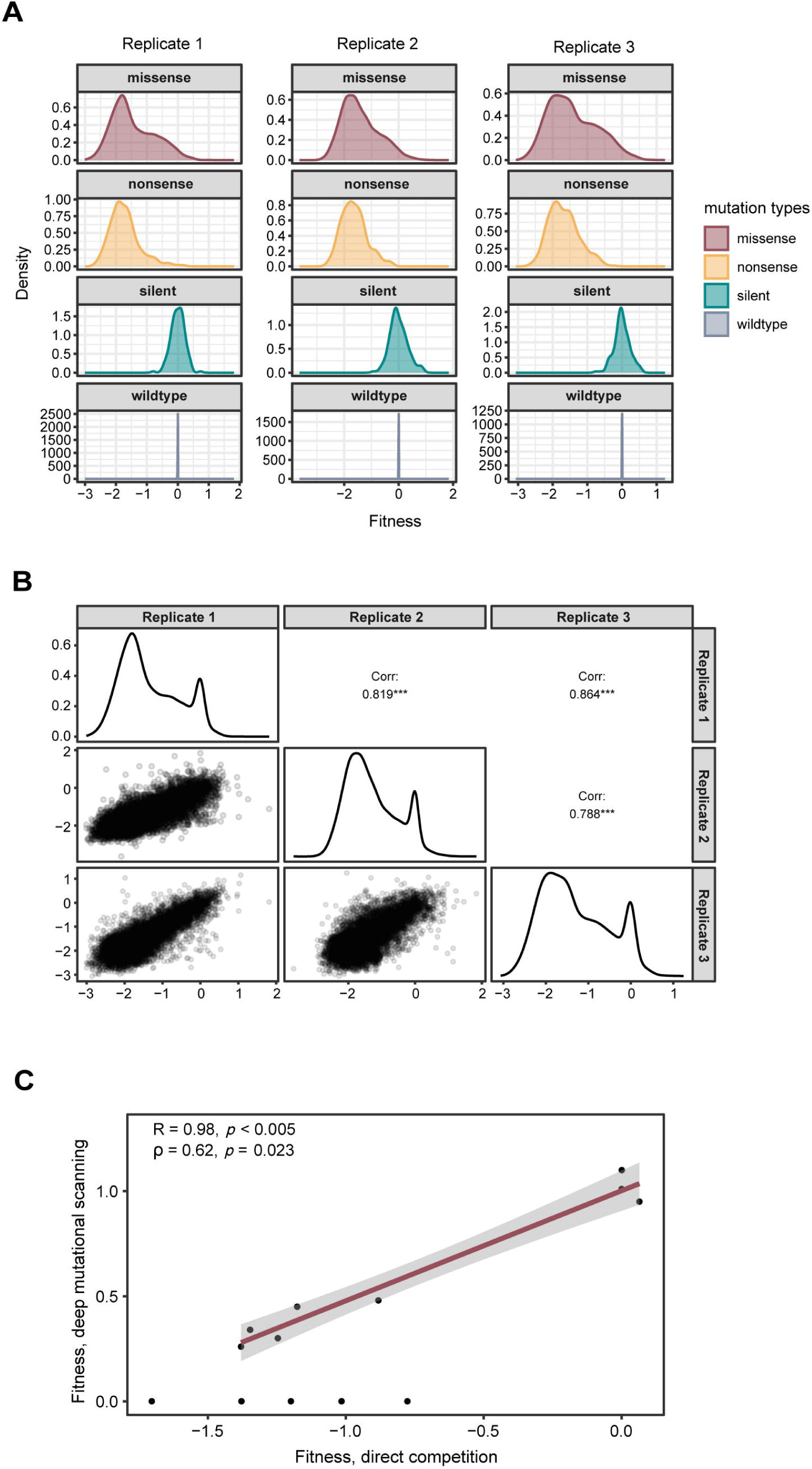
Precision and accuracy of replicative fitness, as measured by deep mutational scanning. (A) The fitness distribution of missense, nonsense, and silent mutations. (B) Correlations of variant fitness in three replicates. The upper right panels show the Pearson correlation coefficients of corresponding replicates with the significance level. Diagonal panels show the overall fitness distribution, disregarding the types of mutation. The lower left panels show the fitness values for individual mutants in the indicated replicates. (C) The fitness values of 16 selected mutations were measured by deep mutational scanning or pairwise competition with the wild type virus. Lethal mutations in the competition assay (fitness = 0) are shown on the x-axis. R indicates the Pearson correlation coefficient among viable variants, while ρ indicates the Spearman correlation coefficient in all variants, including the lethal mutations. The red line shows the trendline using a linear regression model. The gray zone indicates the 95% confidence interval for predictions from the linear model.

We validated our fitness measurements by comparing the deep mutational scanning fitness of 16 amino acid substitutions to the fitness values we’ve measured previously by pairwise competition and quantitative RT-PCR (46). These 16 PB1 substitutions were measured in the same genomic background (A/WSN/33/H1N1) with pairwise competition assays performed in the same cells (A549), at the same MOI (0.01), and for the same number of passages (four). The fitness values in two experiments were well correlated with a Pearson correlation coefficient of 0.98 (*p* < 0.005) for viable variants and a Spearman correlation coefficient of 0.62 (*p* = 0.023) for all variants including the lethal mutants (Figure 4C, Supplemental Figure 2A, B). One substitution (E519D) was lethal in the targeted mutagenesis and not observed in deep mutational scanning. Five other lethal substitutions were identified in passaged DMS libraries, but with very low fitness values.

### Site entropy defines constraints

We calculated site entropy, or Shannon entropy at each site, based on the enrichment of all amino acid variants at a site. The enrichment of each amino acid variant in the calculation was determined by its enrichment ratio after four passages and normalized to sum to 1 (see Methods). High site entropy indicates that variation at the amino acid level does not substantially impact viral fitness and/or that several amino acids are equally tolerated at a site. Because the site entropy calculation would be misleading if some amino acids were absent in the initial libraries, we marked and excluded 16 sites with fewer than 40% of amino acid variants (fewer than 9 out of 21 possible variants) generated in the plasmid libraries (Figure 5, Supplemental Figure 3).

**Figure 5.**
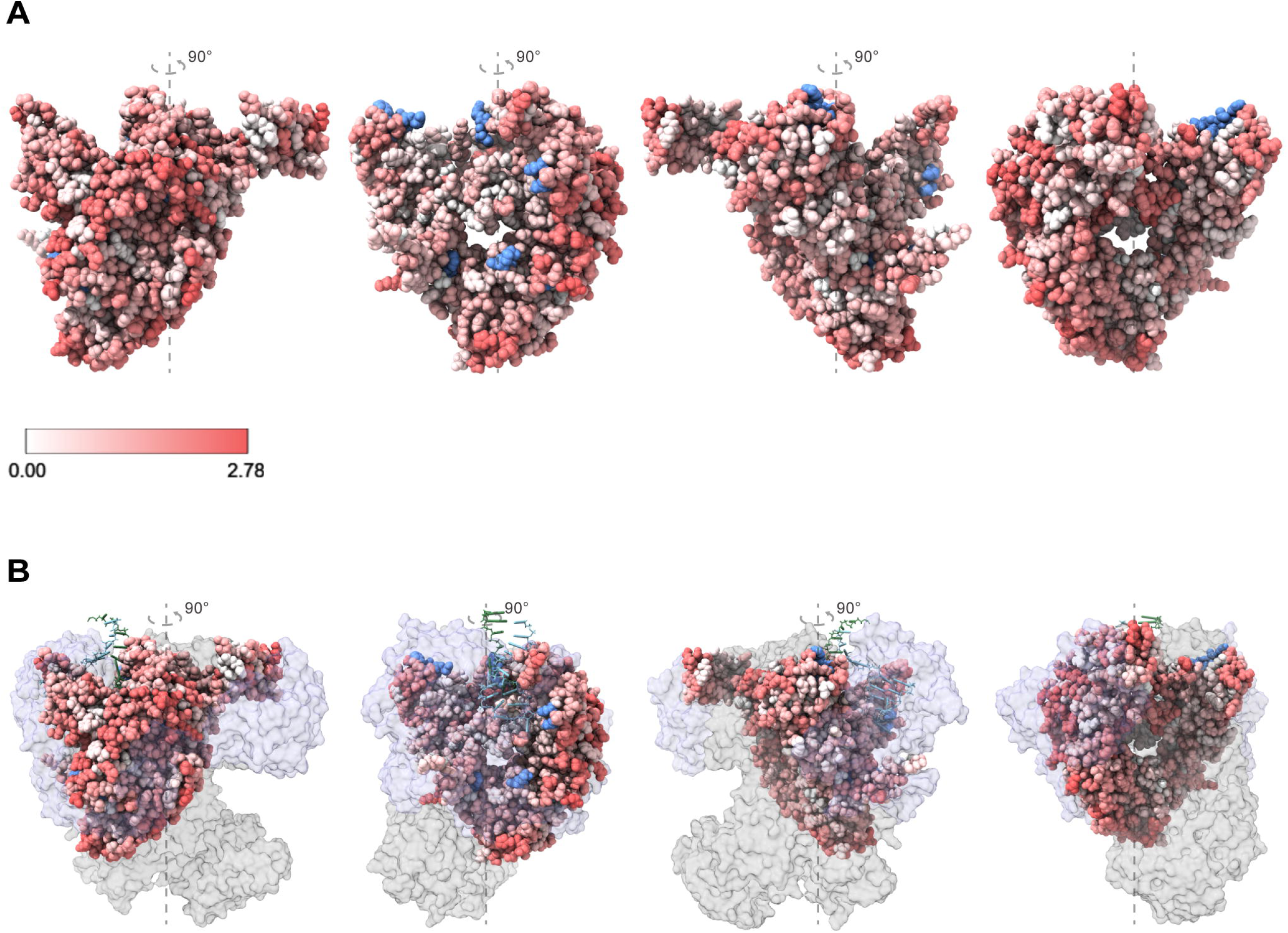
Site entropy of all residues on influenza H1N1 RdRp. (A) Views of site entropy mapped on the H1N1 A/WSN/1933 PB1 structure (PDB: 7NHX). Each view turns 90 degrees counterclockwise from the previous with a fixed y-axis. The coloring ranges from white (lowest, 0) to red (highest, 2.78). Blue indicates the residues at which site entropy may not be accurate, as fewer than 9 amino acid variants, or 40% of all possible amino acid variants, remained in the plasmid libraries after filtering. (B) Position of PB1 relative to PA (purple), PB2 (gray), 31 vRNA (green), and 51 vRNA (cyan). While PB1 is colored by site entropy as that in (A), PA and PB2 are shown by semi-transparent surfaces, and RNAs are presented as ladders.

Site entropy varied across PB1 subdomains. Structural mapping revealed lower site entropy at buried sites and at interfaces between PB1 and RNA and between PB1 and either PA or PB2 (Figure 5A, B). Consistent with this observation, there was a modest, but statistically significant correlation (ρ = 0.28, *p* < 0.005) between site entropy and a residue’s Accessible Surface Area (ASA) (Supplemental Figure 4A). Residues that are more flexible are often more tolerant to mutation and evolve at a higher rate (19). We performed a molecular dynamics simulation and found the correlation between residue flexibility, captured by the Root Mean Square Fluctuation (RMSF) of a 20 ns molecular dynamics simulation of A/Brevig Mission/1/1918(H1N1) RdRp, and site entropy was also weak but significant (ρ = 0.21, *p* < 0.005) (Supplemental Figure 4B). Using the subdomains defined in (42), we grouped site entropy by subdomain. Residues in the fingertips subdomain exhibited lower entropy (p < 0.005 compared to β-hairpin, C-terminal, fingers, prime loop, and ribbon subdomains; p < 0.05 compared to palm and thumb subdomains, p > 0.05 compared to N-terminal subdomain), residues in the prime loop and ribbon subdomains exhibited higher entropy (for prime loop subdomain: p < 0.005 compared to fingertips, N-terminal, Palm, and thumb subdomains, p < 0.5 compared to C-terminal and fingers subdomains; for ribbon subdomain: p < 0.005 compared to fingertips and thumb subdomains; p < 0.5 compared to N-terminal and palm subdomains), and the distribution of entropy values across other subdomains were largely similar (Supplemental Figure 4C).

Because ASA, RMSF, and simple subdomain identity may mask important differences by averaging over a number of high and low entropy sites, we focused subsequent analyses on specific sites with defined functions. The PB1 active site consists of the evolutionarily conserved motifs A-G, with the catalytic metal ions being coordinated by motif A and C at the edge of the central cavity (25, 47). We used logo plots to display the enrichment of each amino acid substitution at residues in motif C (48). The site entropy for the active site was quite low, and there were few alternatives to the wild type amino acid (Figure 6A). Similarly, we evaluated PB1 residues that bound to RNA by hydrogen bonds or interact with RNA due to proximity at each stage (e.g., apo-enzyme, early template binding, late template binding, cap-snatching, pre-initiation, mixed-initiation, initiation-to-catalysis, catalysis-to-nucleotide-incorporation, elongation, and termination). Here, residues interacting with the template RNA (31 vRNA) and product (mRNA) had lower site entropy than others. Site entropy at residues that bind to RNA (5⍰ vRNA) but that are not involved in transcription, were not significantly different than other sites (Figure 6B).

**Figure 6.**
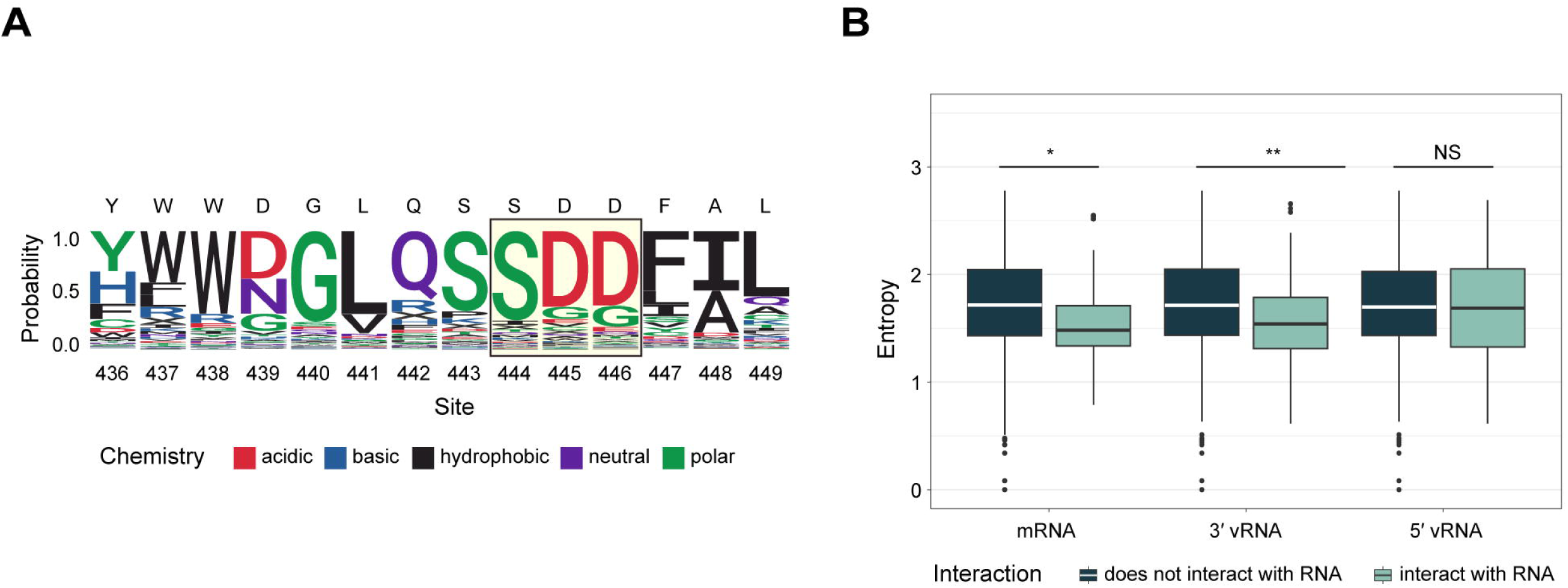
Site entropy of key residues. (A) Enrichment of amino acid substitutions at each residue in motif C. Residues conserved in all negative sense RNA viruses are marked with the light-yellow box. Amino acids are colored by their biochemical characteristics. A stop codon is represented by “X”. (B) Site entropy of sites based on their direct interaction with mRNA, 31 vRNA, and 51 vRNA, visualized by Tukey boxplot. The line in the boxes represent the median, and the top and bottom of the boxes represent 25^th^ and 75^th^ percentile. Data points greater than 75^th^ percentile + 1.5 × interquartile range (IQR) or less than 25^th^ percentile – 1.5 × IQR are shown outside the box and the whisker. Wilcoxon test. * p < 0.5; ** p < 0.05; NS: non-significant.

### Beneficial residues observed in natural evolution

To gain insights into the relationship between mutational tolerance and the long-term evolution of PB1, we compared our measured site entropy to the “natural” amino acid Shannon diversity of each site. The calculation of Shannon diversity in natural sequences is slightly different from that of site entropy for deep mutational scanning; they use the same equation (see Methods), but the former uses the frequency of each amino acid variant, while the latter uses enrichment ratio. We divided the records of naturally evolved PB1 sequences available on GISAID into pre- and post-2009 subsets, separated by the time period of 2009 H1N1 pandemic, to minimize the impact of co-circulation of pre- and post-pandemic viruses. After filtering, we evaluated 1,491 PB1 sequences in the pre-2009 dataset and 35,501 in the post-2009 dataset. Since Shannon diversity is biased for mutations observed in years that have been sampled more densely, we corrected for the uneven sampling of PB1 sequences over time by calculating weighted Shannon diversity as previously described (42, see Methods). In general, there was greater Shannon diversity in the pre-2009 dataset (Supplemental Figure 5). We found a moderate correlation between DMS site entropy and natural Shannon diversity in the pre- and post-2009 datasets (pre- 2009: ρ = 0.40, *p* < 0.005; post-2009: ρ = 0.31, *p* < 0.005. Figures 7A, B).

**Figure 7.**
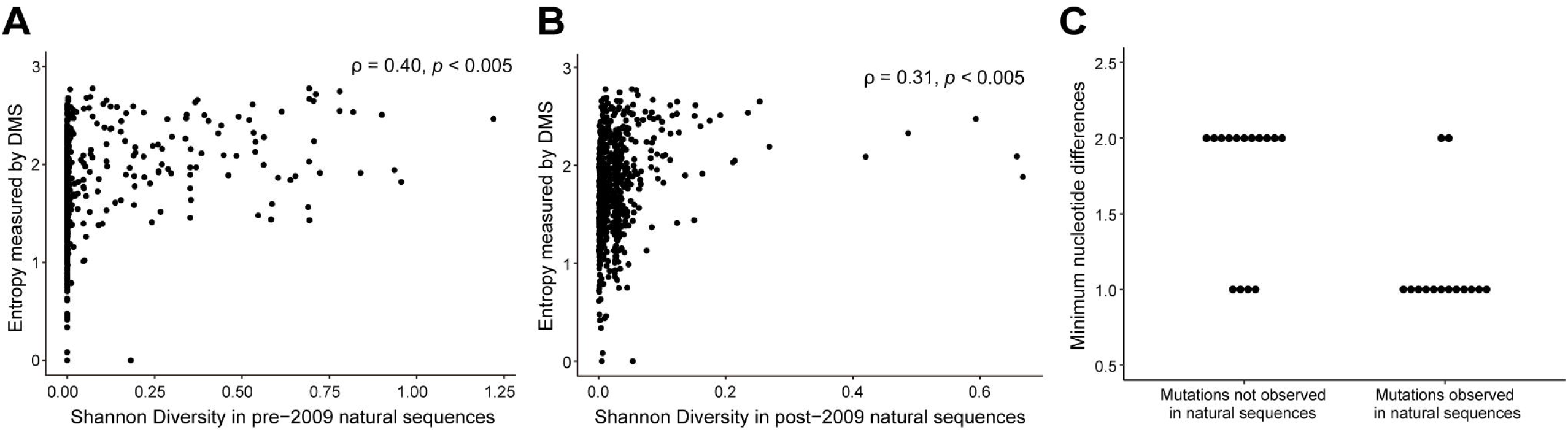
Indications of DMS fitness and mutational tolerance on natural PB1 evolution. Correlation between the Shannon diversity of naturally occurring sequences (A) before and (B) after 2009 and the site entropy measured by deep mutational scanning. ρ indicates the Spearman correlation coefficient. (C) The minimum nucleotide differences between the wild type (or dominant amino acid) in naturally occurring PB1 sequences and beneficial mutations identified by deep mutational scanning. Each dot represents a beneficial mutation.

Similarly, we determined whether mutations identified as beneficial in deep mutational scanning forecast those that appear in the natural evolution of PB1. We defined 29 mutations as beneficial based on a measured fitness greater than two standard deviations (z-score > 2) above the mean fitness of silent mutations (the neutral, null model) (Table 2). All beneficial amino acid mutations had one or two nucleotide changes compared to the corresponding wild type codon. Fourteen of 29 beneficial mutations have occurred during the evolution of H1N1 PB1, and of these, many have appeared multiple times independently. The other 15 were not observed in the available sequences. The majority of beneficial mutations that did appear in natural evolution are accessible by a single nucleotide substitution from the wild type codon, while those that did not appear in natural sequences usually required two nucleotide substitutions in the codon (Figure 7C). Of the beneficial mutations, M317V, T323M, I637V, K653R, K691R, and M744A have appeared in > 0.1% of all sequences collected in at least one year when the mutation was present. Notably, although 691K was the wild type in WSN33 at site 691, arginine (R) was the dominant amino acid in pre-2009 strains (Supplemental Figure 7), suggesting a true fitness advantage of arginine over lysine at this site. Lysine was again the dominant amino acid in the 2009 pandemic strain, but 691R has been detected every year.

**Table 2.**
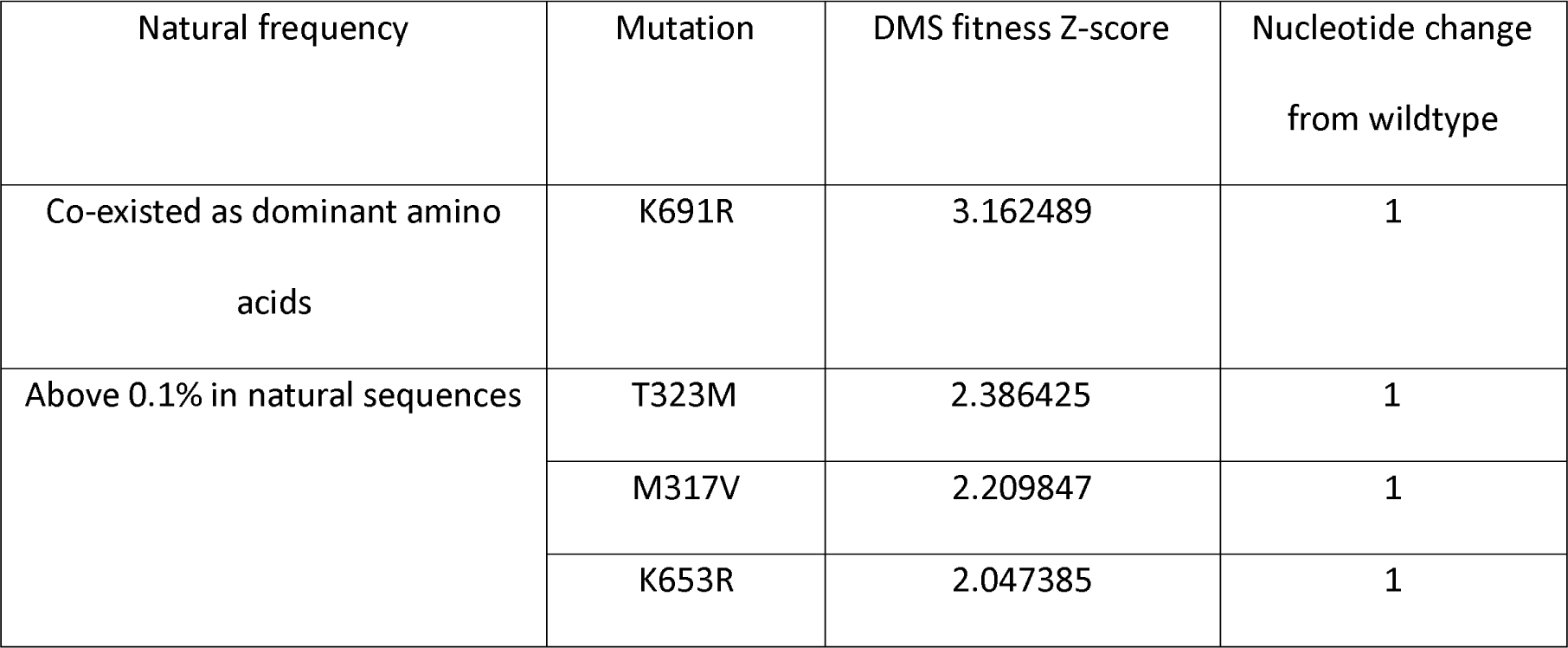
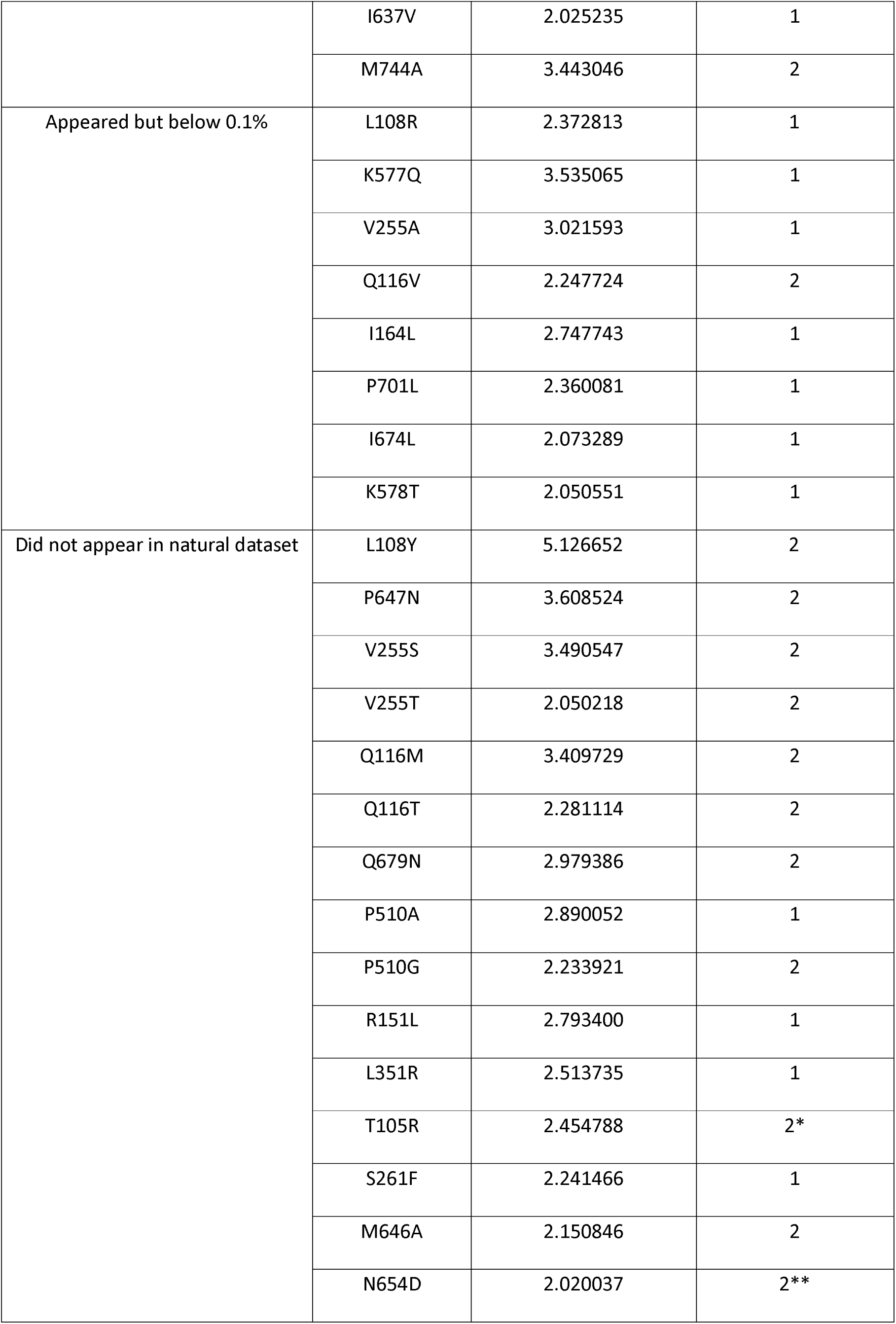
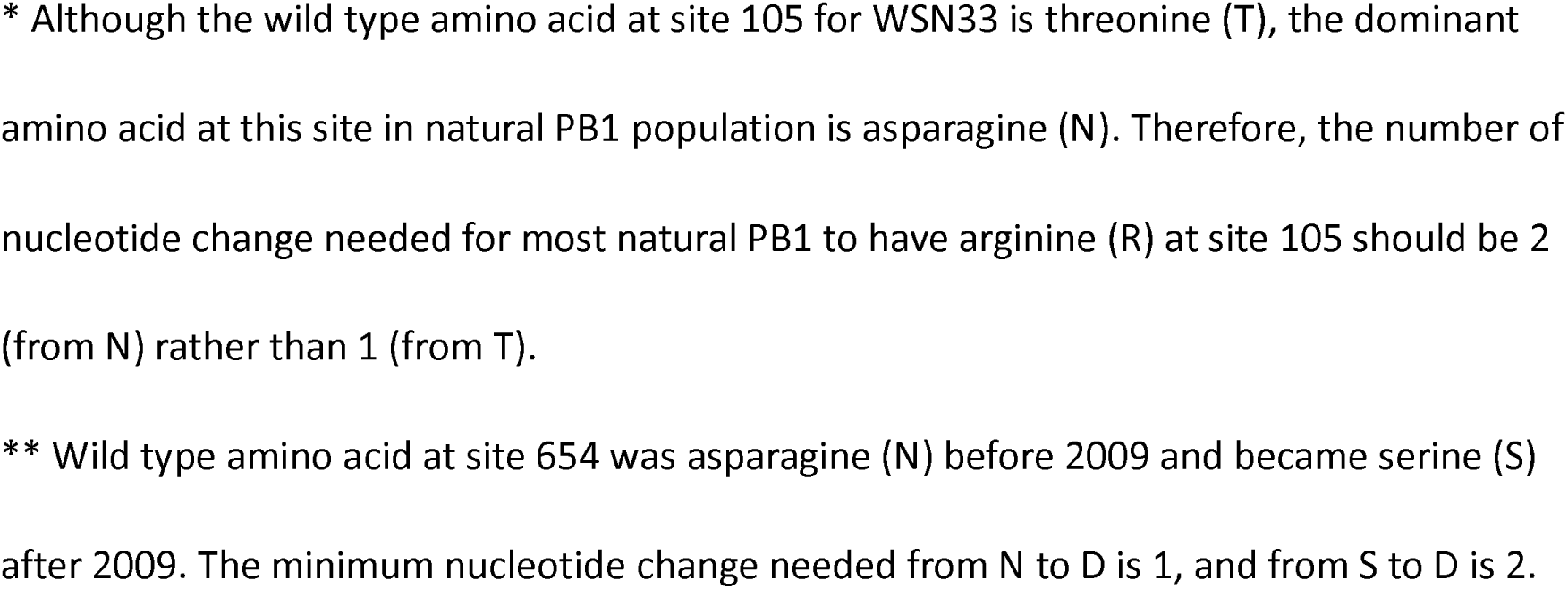
Natural occurrence of beneficial mutations identified by deep mutational scanning.

| Natural frequency | Mutation | DMS fitness Z-score | Nucleotide change<br>from wildtype |
| --- | --- | --- | --- |
| Co-existed as dominant amino<br>acids | K691R | 3.162489 | 1 |
| Above 0.1% in natural sequences | T323M | 2.386425 | 1 |
|  | M317V | 2.209847 | 1 |
|  | K653R | 2.047385 | 1 |
|  | I637V | 2.025235 | 1 |
|  | M744A | 3.443046 | 2 |
| Appeared but below 0.1% | L108R | 2.372813 | 1 |
|  | K577Q | 3.535065 | 1 |
|  | V255A | 3.021593 | 1 |
|  | Q116V | 2.247724 | 2 |
|  | I164L | 2.747743 | 1 |
|  | P701L | 2.360081 | 1 |
|  | I674L | 2.073289 | 1 |
|  | K578T | 2.050551 | 1 |
| Did not appear in natural dataset | L108Y | 5.126652 | 2 |
|  | P647N | 3.608524 | 2 |
|  | V255S | 3.490547 | 2 |
|  | V255T | 2.050218 | 2 |
|  | Q116M | 3.409729 | 2 |
|  | Q116T | 2.281114 | 2 |
|  | Q679N | 2.979386 | 2 |
|  | P510A | 2.890052 | 1 |
|  | P510G | 2.233921 | 2 |
|  | R151L | 2.793400 | 1 |
|  | L351R | 2.513735 | 1 |
|  | T105R | 2.454788 | 2* |
|  | S261F | 2.241466 | 1 |
|  | M646A | 2.150846 | 2 |
|  | N654D | 2.020037 | 2** |

\* Although the wild type amino acid at site 105 for WSN33 is threonine (T), the dominant amino acid at this site in natural PB1 population is asparagine (N). Therefore, the number of nucleotide change needed for most natural PB1 to have arginine (R) at site 105 should be 2 (from N) rather than 1 (from T).
\*\* Wild type amino acid at site 654 was asparagine (N) before 2009 and became serine (S) after 2009. The minimum nucleotide change needed from N to D is 1, and from S to D is 2.

Additional mutations that we identified as beneficial have also been found to be relevant to polymerase activity and viral fitness. Mutations at site 317 were identified in the 1997 Hong Kong H5N1 outbreak (49) and were found to be functionally significant for virulence in mammals (50, 51). Site 744 is located in the vRNA-binding region, and M744V was found to be a canine-adaptive mutation of avian H3N2 (52). Site 674 is both part of the contact points between the PB1 C-terminal and PB2 N-terminal subdomains and interacts with the 31 end of the vRNA promoter; mutations to T, L, and S all increased polymerase activity (53). At the polymerase dimer interface, residue 577 interacts with PA and PB2 (54, 55), and residue 578 orients to a residue in the PB2 unstructured loop; K577E in avian H9N2 increases polymerase activity at a lower replication temperature (56) and serial passage of A/Hong Kong/1/68 (H3N2) in mice also gave rise to K577E/M/Q (57). Lysine 578, the wild type, is a ubiquitination site, and mutations from K578 to both non-charged alanine (A) and positively charged arginine (R) increase polymerase activity but are harmful to viral fitness (58). The neutral side chain of A578 reduced polymerase dimerization, while the positively charged R578 aborted cRNA synthesis and led to the premature assembly of the dimer.

## Discussion

We performed a near complete deep mutational scan of the WSN33 PB1 RdRp subunit, defining the impacts of nearly all amino acid substitutions on replicative fitness. Most substitutions are detrimental, and we identified mutational constraints at sites involved in key polymerase interactions, specifically at sites interacting with the RNA template and product. In contrast, mutations in other regions of the protein are better tolerated. Diversity at these sites was moderately correlated with site diversity as defined in available influenza sequences. A small number of mutations are beneficial, and many of these have been observed in natural evolution. Those that were not observed in natural evolution were generally inaccessible by single nucleotide mutation. Our study was comprehensive, as we interrogated a much larger number of codon and amino acid variants compared to studies that evaluate mutations occurring in natural sequences or generated by error-prone PCR. While prior work on the functional domains and evolutionary constraints on RdRp have largely relied on analyses of sequence conservation, our DMS identified significant, site-specific heterogeneity in the influenza virus polymerase (59).

Unlike in hemagglutinin (60) and neuraminidase (31), we find that the evolutionary constraints on the influenza virus RdRp are not well defined by protein subdomain. Instead, each subdomain has some sites that are under strict purifying selection and other sites that are more tolerant to mutation. Similar phenomena were observed in naturally occurring genomes, where conservative and variable residues were distributed relatively evenly across major subdomains (61). These findings highlight the importance of local structures and functional interactions in influenza virus replication. As expected, mutations to amino acids with side chains of similar biochemical properties (e.g., charged/uncharged, polar/non-polar) are usually more tolerated. This is consistent with the impact of these biochemical properties on higher-level protein structures: large and non-polar amino acids are more likely to form hydrophobic cores, while small and charged amino acids are intrinsically disordered (62).

We identified beneficial mutations that have been observed in the natural evolution of influenza virus RdRp and found accessibility by single nucleotide substitution to be a key factor determining whether a beneficial mutation can arise naturally. We also identified several adaptative mutations that arose in nature with more than one nucleotide change, which could imply an indirect evolutionary path involving gain and subsequent loss of intermediate mutations (63). Many of the beneficial mutations identified in our study not only increase polymerase activity but have been shown to be functionally important for host adaptation or by altering post-translation modification. In addition, mutations with higher fitness had a moderate but significant association with sites that have higher mutational tolerance.

Our work is subject to several limitations. First, while our deep mutational scan provides comprehensive fitness measurements in the WSN33 genetic background, the measured mutational effects may not be recapitulated in the genetic background of other H1N1 strains. Second, we performed our DMS on A549 cells, which allow for high volume infections and the robust viral replication necessary for a comprehensive screen with a large library. It is possible that fitness values may differ in a more physiologically relevant replication system, such as primary airway epithelial cells. Third, we focused on the mutational effects of single amino acid substitutions and did not account for epistatic interactions within PB1 and between PB1 and other viral proteins. Finally, we only examined fitness in terms of replication, but various treatments can be applied to the variant virus library and future research can examine mutational fitness under specific conditions such as with drug selection or altered baseline mutational rates.

Overall, we have developed a comprehensive map of the local fitness landscape for the influenza A virus PB1 protein. In doing so, we identified how specific amino acid substitutions affect the replicative fitness of the virus and the degree of evolutionary constraint at each site. Our work provides a foundation for subsequent studies of influenza virus replication and host adaptation and may prove to be a valuable addition to genomic surveillance efforts.

## Acknowledgements

We thank Jesse Bloom and Shirleen Soh for making their analysis code available and for helpful suggestions, and Aaron King, Gideon Bradburd, and Kayla Peck for helpful discussion. We further acknowledge the contributions of all submitters to GISAID. We performed molecular graphics with UCSF ChimeraX, developed by the Resource for Biocomputing, Visualization, and Informatics at the University of California, San Francisco, with support from National Institutes of Health R01-GM129325 and the Office of Cyber Infrastructure and Computational Biology, National Institute of Allergy and Infectious Diseases. The MD simulations were performed on computational resources managed and supported by Princeton Research Computing, a consortium of groups including the Princeton Institute for Computational Science and Engineering (PICSciE) and the Office of Information Technology’s High Performance Computing Center and Visualization Laboratory at Princeton University. This work was supported by NIH R01 AI170520 and a Burroughs Wellcome Fund Investigator in the Pathogenesis of Infectious Diseases Award, both to ASL, and NIH DP2 AI175474 to ATV.

## Supplemental Figure Legends

**Supplemental Figure 1. Full description of deep mutational scanning libraries.** (A) Raw sequencing reads of each sample. The “filter” in “fail filter” refers to general Illumina filters. “Low Q barcode” refers to sequences having any nucleotide with a Q-score below 15 in the 16× N molecule-specific barcodes. Sequences that failed the filter or with low-Q barcodes were discarded in subsequent analyses. (B) The number of distinct barcodes observed in each sample. Each barcode needs to be observed at least twice to determine the consensus sequence for that contig. The bar at 1 corresponds either to barcodes that were only observed once or to sequencing errors that gave rise to new barcodes. (C) Barcodes after aligning to wild type WSN33 PB1 sequence. “Too few reads” corresponds to the bar at 1 in panel (B). Sequences categorized as “too few reads” were removed from subsequent analyses. (D) Sequencing depth at each site in PB1 after removing contigs with too few reads. The number of counts includes the codon counts for both variant and wild type codons. (E) The mutational frequency at each site in PB1 after removing contigs with too few reads. The spike at site 577 in library Rep1P0, Rep1P4, Rep3P0, and Rep3P4 is likely an issue with the sequencing library preparation, as other sequencing runs using the same samples did not show such peaks (data not shown). The spikes have little impact on the type of codons present in the libraries. The impact of peaks on fitness measurements is also minimal since we compared the passaged libraries to the plasmid libraries. (F) Mutation sampling completeness. The plot shows the fraction of codon and amino acid mutations observed no more than the indicated number of times. This plot describes both variant diversity and sequencing completeness in a library. (G) Frequency of different types of nucleotide change. The plot shows nucleotide change among mutations with only one nucleotide change and works as a check for oxidative damage. An excessive number of C to A or G to T mutations suggests potential oxidative damage. The plot shows no over-representation of either mutation in the libraries.

**Supplemental Figure 2. Fitness comparison between deep mutational scanning and direct competition in early passages.** The comparison between the replicative fitness measured by direct competition with the wild type strain and by deep mutational scanning (A) After virus library rescue, before passaging, and (B) after one passage on A549 cells. R indicates the Pearson correlation coefficient for viable variants, while ρ indicates the Spearman correlation coefficient for all variants including lethal mutations. The red line shows the trendline using a linear regression model. The gray zone indicates the 95% confidence interval for predictions from the linear model.

**Supplemental Figure 3. Sites with varying mutational representation.** Count of sites with different percentages of amino acid variants present at that site in the plasmid libraries, after filtering out the mutations with low sequencing counts or with sequencing library preparation errors. One hundred percent indicates that twenty amino acid variants plus variants for stop codons were all observed at a site.

**Supplemental Figure 4. Correlation between site entropy and defined features on RdRp.** Correlation between a residue’s (A) accessible surface area or (B) root mean square fluctuation, and its site entropy. Each dot represents a residue on the RdRp. ρ indicates the Spearman correlation coefficient. (C) Site entropy distribution in different subdomains of RdRp. The chart below shows the adjusted p-values by Bonferroni correction between each pair of subdomain comparisons.

**Supplemental Figure 5. Amino acid diversity at sites of naturally occurring influenza H1N1 PB1 sequences.** Weighted Shannon diversity for each site in natural PB1 evolution. Diversity in pre- and post-2009 sequences was calculated separately.

**Supplemental Figure 6. Correlation between site entropy and mutational fitness.** Each dot represents an amino acid substitution at a site. ρ indicates the Spearman correlation coefficient.

**Supplemental Figure 7. Frequency change of amino acid variants at site 691.** Frequency of amino acid variants observed at site 691 of naturally occurring PB1 sequences from 1934 to 2023. “X” stands for uncertain/ambiguous amino acid. Dominant amino acid variants were labeled on the plot.

## References

1. Zhang X, Li Y, Jin S, Zhang Y, Sun L, Hu X, Zhao M, Li F, Wang T, Sun W, Feng N, Wang H, He H, Zhao Y, Yang S, Xia X, Gao Y. 2021. PB1 S524G mutation of wild bird-origin H3N8 influenza A virus enhances virulence and fitness for transmission in mammals. Emerging Microbes & Infections 10:1038–1051.

2. Feng X, Wang Z, Shi J, Deng G, Kong H, Tao S, Li C, Liu L, Guan Y, Chen H. 2016. Glycine at Position 622 in PB1 Contributes to the Virulence of H5N1 Avian Influenza Virus in Mice. Journal of Virology 90:1872–1879.

3. Xu C, Hu W-B, Xu K, He Y-X, Wang T-Y, Chen Z, Li T-X, Liu J-H, Buchy P, Sun B. 2012. Amino acids 473V and 598P of PB1 from an avian-origin influenza A virus contribute to polymerase activity, especially in mammalian cells. Journal of General Virology 93:531–540.

4. Goldhill DH, te Velthuis AJW, Fletcher RA, Langat P, Zambon M, Lackenby A, Barclay WS. 2018. The mechanism of resistance to favipiravir in influenza. Proceedings of the National Academy of Sciences 115:11613–11618.

5. Li J, Liang L, Jiang L, Wang Q, Wen X, Zhao Y, Cui P, Zhang Y, Wang G, Li Q, Deng G, Shi J, Tian G, Zeng X, Jiang Y, Liu L, Chen H, Li C. 2021. Viral RNA-binding ability conferred by SUMOylation at PB1 K612 of influenza A virus is essential for viral pathogenesis and transmission. PLOS Pathogens 17:e1009336.

6. Varga ZT, Ramos I, Hai R, Schmolke M, García-Sastre A, Fernandez-Sesma A, Palese P. 2011. The Influenza Virus Protein PB1-F2 Inhibits the Induction of Type I Interferon at the Level of the MAVS Adaptor Protein. PLoS Pathogens 7:e1002067.

7. Hai R, Schmolke M, Varga ZT, Manicassamy B, Wang TT, Belser JA, Pearce MB, García- SastreA, Tumpey TM, Palese P. 2010. PB1-F2 Expression by the 2009 Pandemic H1N1 Influenza Virus Has Minimal Impact on Virulence in Animal Models. Journal of Virology 84:4442–4450.

8. Laporte M, Stevaert A, Raeymaekers V, Boogaerts T, Nehlmeier I, Chiu W, Benkheil M, Vanaudenaerde B, Pöhlmann S, Naesens L. 2019. Hemagglutinin Cleavability, Acid Stability, and Temperature Dependence Optimize Influenza B Virus for Replication in Human Airways. Journal of Virology 94.

9. Goñi N, Iriarte A, Comas V, Martín Soñora, Moreno P, Moratorio G, Musto H, Cristina J. 2012. Pandemic influenza A virus codon usage revisited: biases, adaptation and implications for vaccine strain development. Virology Journal 9.

10. Fan RZ, Eric, Chloe, Olive, Nicholls JM, Rabadan R, Peiris M, Leo L.M. Poon. 2015. Generation of Live Attenuated Influenza Virus by Using Codon Usage Bias. Journal of Virology 89:10762–10773.

11. Kumar N, Bera BC, Greenbaum BD, Bhatia S, Sood R, Selvaraj P, Anand T, Tripathi BN, Virmani N. 2016. Revelation of Influencing Factors in Overall Codon Usage Bias of Equine Influenza Viruses. PLOS ONE 11:e0154376.

12. Mintseris J, Weng Z. 2005. Structure, function, and evolution of transient and obligate protein-protein interactions. Proceedings of the National Academy of Sciences 102:10930– 10935.

13. Aharoni A, Gaidukov L, Khersonsky O, Gould SM, Roodveldt C, Tawfik DS. 2005. The “evolvability” of promiscuous protein functions. Nature Genetics 37:73–76.

14. Andreeva A, Murzin AG. 2006. Evolution of protein fold in the presence of functional constraints. Current Opinion in Structural Biology 16:399–408.

15. Hom N, Gentles L, Bloom JD, Lee KK. 2019. Deep Mutational Scan of the Highly Conserved Influenza A Virus M1 Matrix Protein Reveals Substantial Intrinsic Mutational Tolerance. Journal of Virology 93:e00161–19.

16. Goldman N, Thorne JL, Jones DT. 1998. Assessing the Impact of Secondary Structure and Solvent Accessibility on Protein Evolution. Genetics 149:445–458.

17. Velázquez-Muriel J, Rueda M, Cuesta I, Pascual-Montano A, Orozco M, José María Carazo. 2009. Comparison of molecular dynamics and superfamily spaces of protein domain deformation. BMC Structural Biology 9.

18. Friedland GD, Nils-Alexander Lakomek, Griesinger C, Meiler J, Kortemme T. 2009. A Correspondence Between Solution-State Dynamics of an Individual Protein and the Sequence and Conformational Diversity of its Family. PLOS Computational Biology 5:e1000393–e1000393.

19. Marsh JA, Teichmann SA. 2013. Parallel dynamics and evolution: Protein conformational fluctuations and assembly reflect evolutionary changes in sequence and structure. BioEssays 36:209–218.

20. Mintseris J, Weng Z. 2005. Structure, function, and evolution of transient and obligate protein-protein interactions. Proceedings of the National Academy of Sciences 102:10930– 10935.

21. Eames M, Kortemme T. 2007. Structural Mapping of Protein Interactions Reveals Differences in Evolutionary Pressures Correlated to mRNA Level and Protein Abundance. Structure 15:1442–1451.

22. Worth CL, Gong S, Blundell TL. 2009. Structural and functional constraints in the evolution of protein families. Nature Reviews Molecular Cell Biology 10:709–720.

23. Franzosa EA, Xia Y. 2009. Structural Determinants of Protein Evolution Are Context-Sensitive at the Residue Level. Molecular Biology and Evolution 26:2387–2395.

24. Ramsey DC, Scherrer MP, Zhou T, Wilke CO. 2011. The Relationship Between Relative Solvent Accessibility and Evolutionary Rate in Protein Evolution. Genetics 188:479–488.

25. te Velthuis AJW, Fodor E. 2016. Influenza virus RNA polymerase: insights into the mechanisms of viral RNA synthesis. Nature Reviews Microbiology 14:479–493.

26. Kouba T, Drncová P, Cusack S. 2019. Structural snapshots of actively transcribing influenza polymerase. Nature Structural & Molecular Biology 26:460–470.

27. York A, Hengrung N, Vreede FT, Huiskonen JT, Fodor E. 2013. Isolation and characterization of the positive-sense replicative intermediate of a negative-strand RNA virus. Proceedings of the National Academy of Sciences of the United States of America 110:E4238–4245.

28. Soh YS, Moncla LH, Eguia R, Bedford T, Bloom JD. 2019. Comprehensive mapping of adaptation of the avian influenza polymerase protein PB2 to humans. eLife 8:e45079.

29. Sourisseau M, Lawrence DA, Schwarz MC, Storrs C, Veit EC, Bloom JD, Evans M. 2019. Deep Mutational Scanning Comprehensively Maps How Zika Envelope Protein Mutations Affect Viral Growth and Antibody Escape. Journal of Virology 93.

30. Starr TN, Greaney AJ, Hilton SK, Ellis D, Crawford KHD, Dingens AS, Navarro MJ, Bowen JE, Tortorici MA, Walls AC, King NP, Veesler D, Bloom JD. 2020. Deep Mutational Scanning of SARS-CoV-2 Receptor Binding Domain Reveals Constraints on Folding and ACE2 Binding. Cell 182:1295–1310.e20.

31. Lei R, Milena A, Tan TC, Qi Wen Teo, Wang Y-Q, Zhang X, Luo S, Nair SK, Peng J, Wu NC. 2023. Mutational fitness landscape of human influenza H3N2 neuraminidase. Cell Reports 42:111951–111951.

32. Doud MB, Hensley SE, Bloom JD. 2017. Complete mapping of viral escape from neutralizing antibodies. PLOS Pathogens 13:e1006271.

33. Bloom JD, Dingens A. 2019. Tiling primers for codon mutagenesis. Github. https://github.com/jbloomlab/CodonTilingPrimers

34. Bloom JD. 2014. An Experimentally Determined Evolutionary Model Dramatically Improves Phylogenetic Fit. Molecular Biology and Evolution 31:1956–1978.

35. Dingens AS, Haddox HK, Overbaugh J, Bloom JD. 2017. Comprehensive Mapping of HIV-1 Escape from a Broadly Neutralizing Antibody. Cell Host & Microbe 21:777–787.e4.

36. Peck K. 2017. RE_check. Github. https://github.com/kmpeck/RE_check

37. Doud M, Bloom J. 2016. Accurate Measurement of the Effects of All Amino-Acid Mutations on Influenza Hemagglutinin. Viruses 8:155.

38. Bloom JD. 2015. Software for the analysis and visualization of deep mutational scanning data. BMC Bioinformatics 16.

39. Centers for Disease Control and Prevention. 2019. 2009 H1N1 Pandemic Timeline. Centers for Disease Control and Prevention. https://www.cdc.gov/flu/pandemic-resources/2009-pandemic-timeline.html.

40. Katoh K. 2002. MAFFT: a novel method for rapid multiple sequence alignment based on fast Fourier transform. Nucleic Acids Research 30:3059–3066.

41. Shannon CE. 1948. A Mathematical Theory of Communication. Bell System Technical Journal 27:379–423.

42. Arcos S, Han AX, te W, Russell CA, Lauring AS. 2023. Mutual information networks reveal evolutionary relationships within the influenza A virus polymerase. Virus Evolution 9.

43. Pettersen EF, Goddard TD, Huang CC, Meng EC, Couch GS, Croll TI, Morris JH, Ferrin TE. 2020. UCSF ChimeraX: Structure visualization for researchers, educators, and developers. Protein Science 30:70–82.

44. Laskowski RA, Swindells MB. 2011. LigPlot+: Multiple Ligand–Protein Interaction Diagrams for Drug Discovery. Journal of Chemical Information and Modeling 51:2778–2786.

45. Krissinel E, Henrick K. 2007. Inference of Macromolecular Assemblies from Crystalline State. Journal of Molecular Biology 372:774–797.

46. Visher E, Whitefield SE, McCrone JT, Fitzsimmons W, Lauring AS. 2016. The Mutational Robustness of Influenza A Virus. PLoS Pathogens 12.

47. Chu C, Fan S, Li C, Macken C, Kim JH, Hatta M, Neumann G, Kawaoka Y. 2012. Functional Analysis of Conserved Motifs in Influenza Virus PB1 Protein. PLoS ONE 7:e36113.

48. Wagih O. 2017. ggseqlogo: a versatile R package for drawing sequence logos. Bioinformatics 33:3645–3647.

49. Katz JM, Lu X, Tumpey TM, Smith CB, Shaw MW, Subbarao K. 2000. Molecular Correlates of Influenza A H5N1 Virus Pathogenesis in Mice. Journal of Virology 74:10807–10810.

50. Lycett SJ, Ward MJ, Lewis FI, Poon AFY, Kosakovsky Pond SL, Brown AJL. 2009. Detection of Mammalian Virulence Determinants in Highly Pathogenic Avian Influenza H5N1 Viruses: Multivariate Analysis of Published Data. Journal of Virology 83:9901–9910.

51. Nao N, Kajihara M, Manzoor R, Maruyama J, Yoshida R, Muramatsu M, Miyamoto H, Igarashi M, Eguchi N, Sato M, Kondoh T, Okamatsu M, Sakoda Y, Kida H, Takada A. 2015. A Single Amino Acid in the M1 Protein Responsible for the Different Pathogenic Potentials of H5N1 Highly Pathogenic Avian Influenza Virus Strains. PLOS ONE 10:e0137989.

52. Li X, Liu J, Qiu Z, Liao Q, Peng Y, Chen Y, Shu Y. 2021. Host-Adaptive Signatures of H3N2 Influenza Virus in Canine. Frontiers in Veterinary Science 8:740472.

53. Welkers MRA, Pawestri HA, Fonville JM, Sampurno OD, Pater M, Holwerda M, Han AX, Russell CA, Jeeninga RE, Setiawaty V, de Jong MD, Eggink D. 2019. Genetic diversity and host adaptation of avian H5N1 influenza viruses during human infection. Emerging Microbes & Infections 8:262–271.

54. Fan H, Walker AP, Carrique L, Keown JR, Serna Martin I, Karia D, Sharps J, Hengrung N, Pardon E, Steyaert J, Grimes JM, Fodor E. 2019. Structures of influenza A virus RNA polymerase offer insight into viral genome replication. Nature 573:287–290.

55. Kuang Yu Chen, Dos E, Enouf V, Isel C, Naffakh N. 2019. Influenza virus polymerase subunits co-evolve to ensure proper levels of dimerization of the heterotrimer. Plos Pathogens 15:e1008034–e1008034.

56. Kamiki H, Matsugo H, Kobayashi T, Ishida H, Takenaka-Uema A, Murakami S, Horimoto T. 2018. A PB1-K577E Mutation in H9N2 Influenza Virus Increases Polymerase Activity and Pathogenicity in Mice. Viruses 10:653.

57. Ping J, Keleta L, Forbes NE, Dankar SK, Stecho W, Tyler S, Zhou Y, Babiuk LA, Weingartl HM, Halpin RA, Boyne A, Bera J, Hostetler J, Fedorova N, Proudfoot K, Katzel DA, Stockwell T, Elodie Ghedin, Spiro DM, Brown EG. 2011. Genomic and Protein Structural Maps of Adaptive Evolution of Human Influenza A Virus to Increased Virulence in the Mouse. PLOS ONE 6:e21740–e21740.

58. Günl F, Krischuns T, Schreiber JA, Henschel L, Wahrenburg M, Drexler HCA, Leidel SA, Cojocaru V, Seebohm G, Mellmann A, Schwemmle M, Ludwig S, Brunotte L. 2023. The ubiquitination landscape of the influenza A virus polymerase. Nature Communications 14:787.

59. Wu NC, Olson CA, Du Y, Le S, Tran K, Remenyi R, Gong D, Al-Mawsawi LQ, Qi H, Wu T-T, Sun R. 2015. Functional Constraint Profiling of a Viral Protein Reveals Discordance of Evolutionary Conservation and Functionality. PLOS Genetics 11:e1005310.

60. Lee JM, Huddleston J, Doud MB, Hooper KA, Wu NC, Bedford T, Bloom JD. 2018. Deep mutational scanning of hemagglutinin helps predict evolutionary fates of human H3N2 influenza variants. Proceedings of the National Academy of Sciences 115.

61. Figueiredo-Nunes I, Trigueiro-Louro J, Rebelo-de-Andrade H. 2023. Exploring new antiviral targets for influenza and COVID-19: Mapping promising hot spots in viral RNA polymerases. Virology 578:45–60.

62. Forman-Kay JD, Mittag T. 2013. From Sequence and Forces to Structure, Function, and Evolution of Intrinsically Disordered Proteins. Structure 21:1492–1499.

63. Wu NC, Dai L, Olson CA, Lloyd-Smith JO, Sun R. 2016. Adaptation in protein fitness landscapes is facilitated by indirect paths. eLife 5:e16965.

