## Supplementary figures and images for "Deep mutational scanning reveals the functional constraints and evolutionary potential of the influenza A virus PB1 protein"

### Supplemental Figure 1

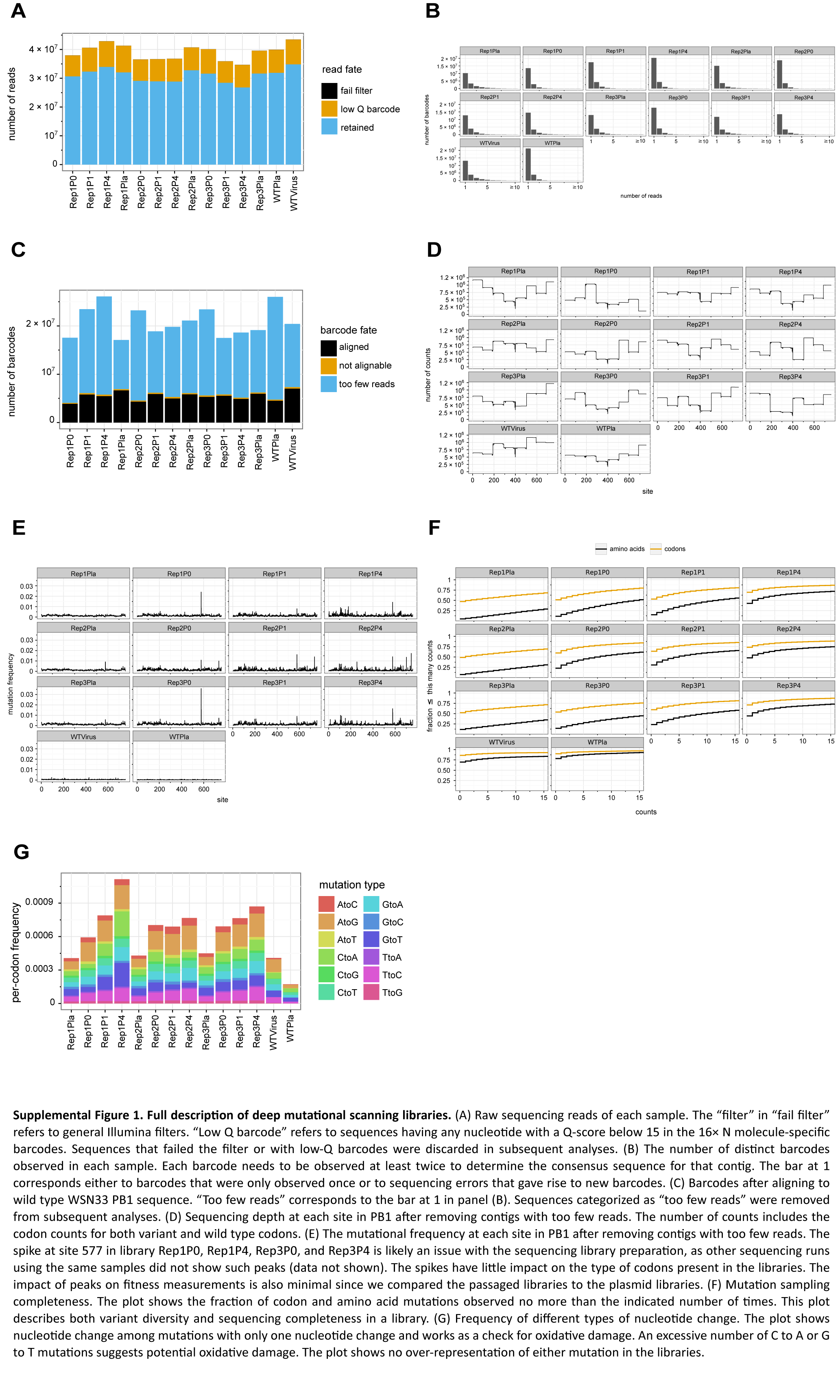

### Supplemental Figure 2

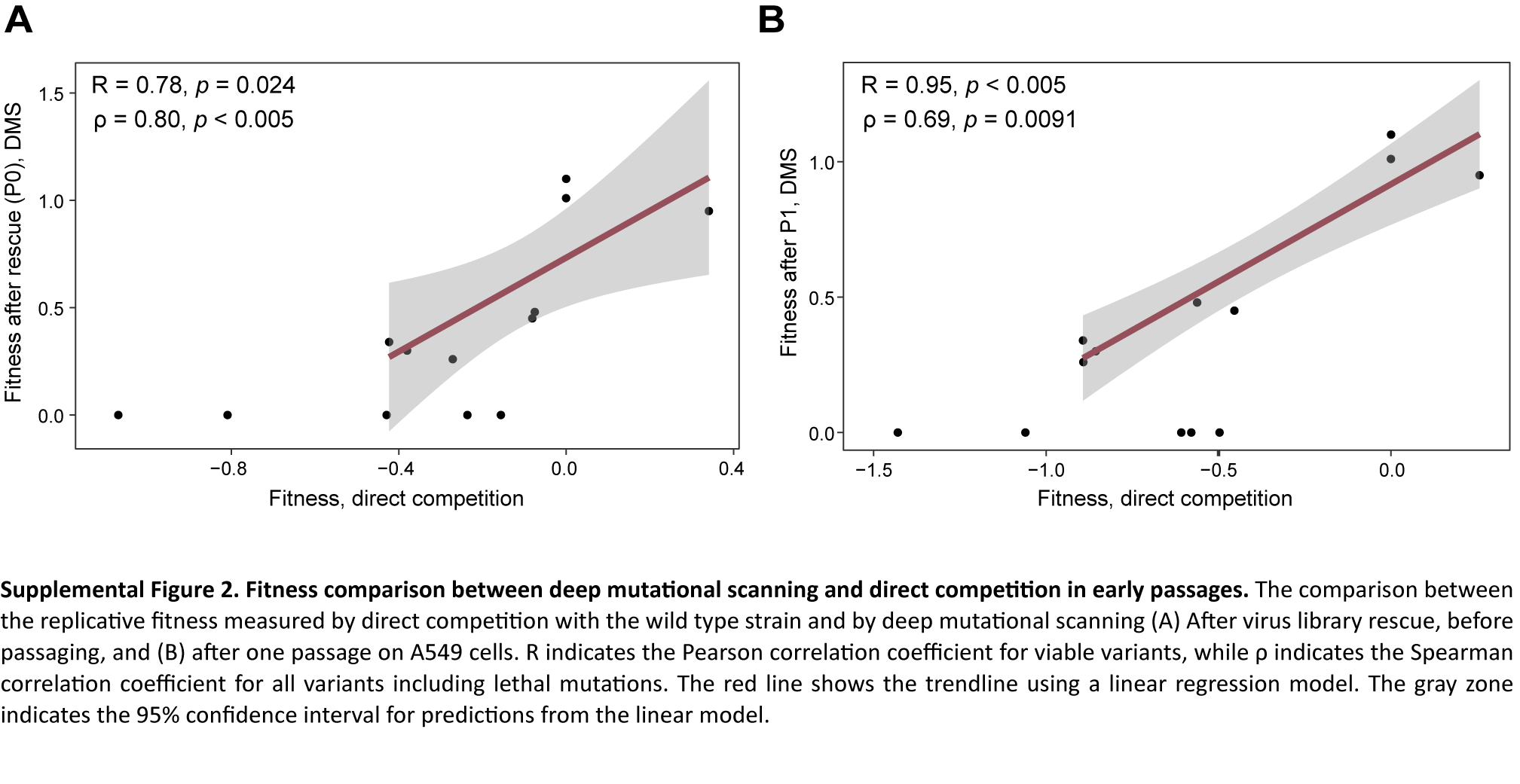

### Supplemental Figure 3

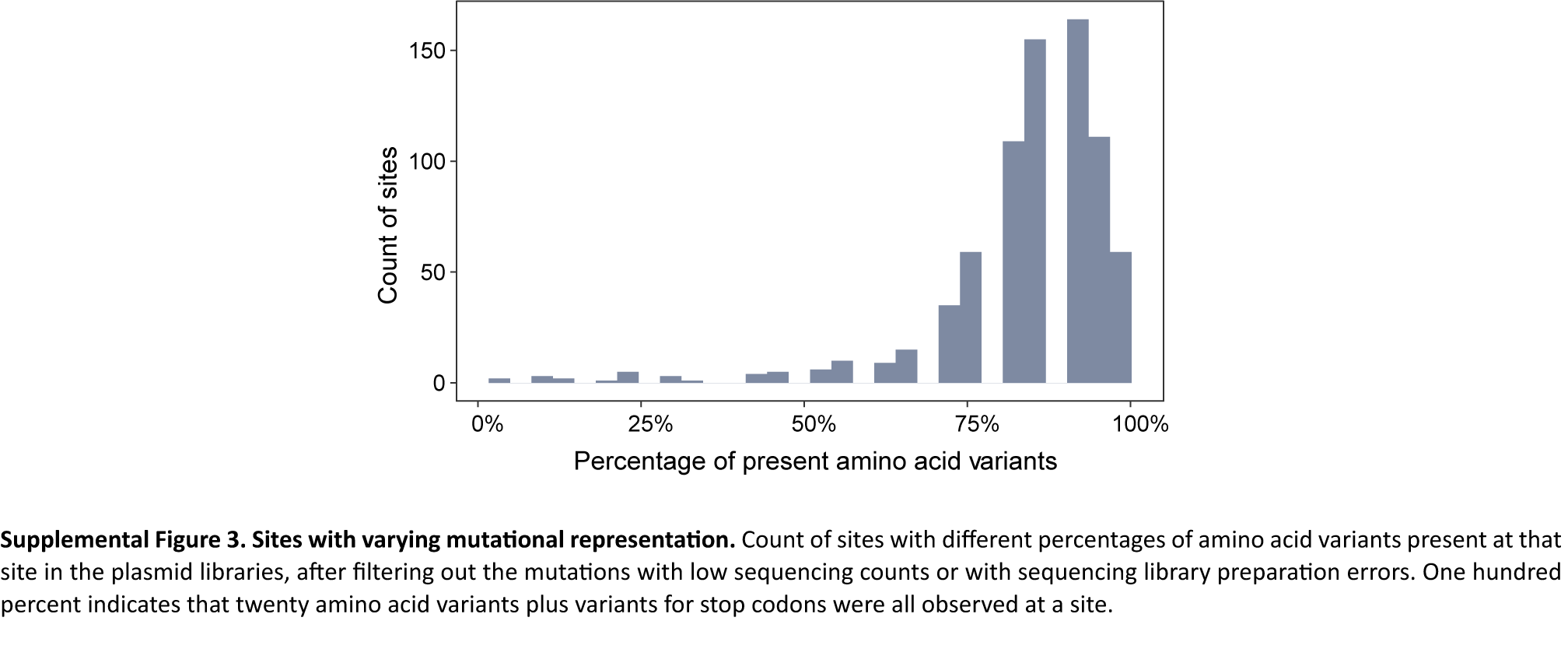

### Supplemental Figure 4

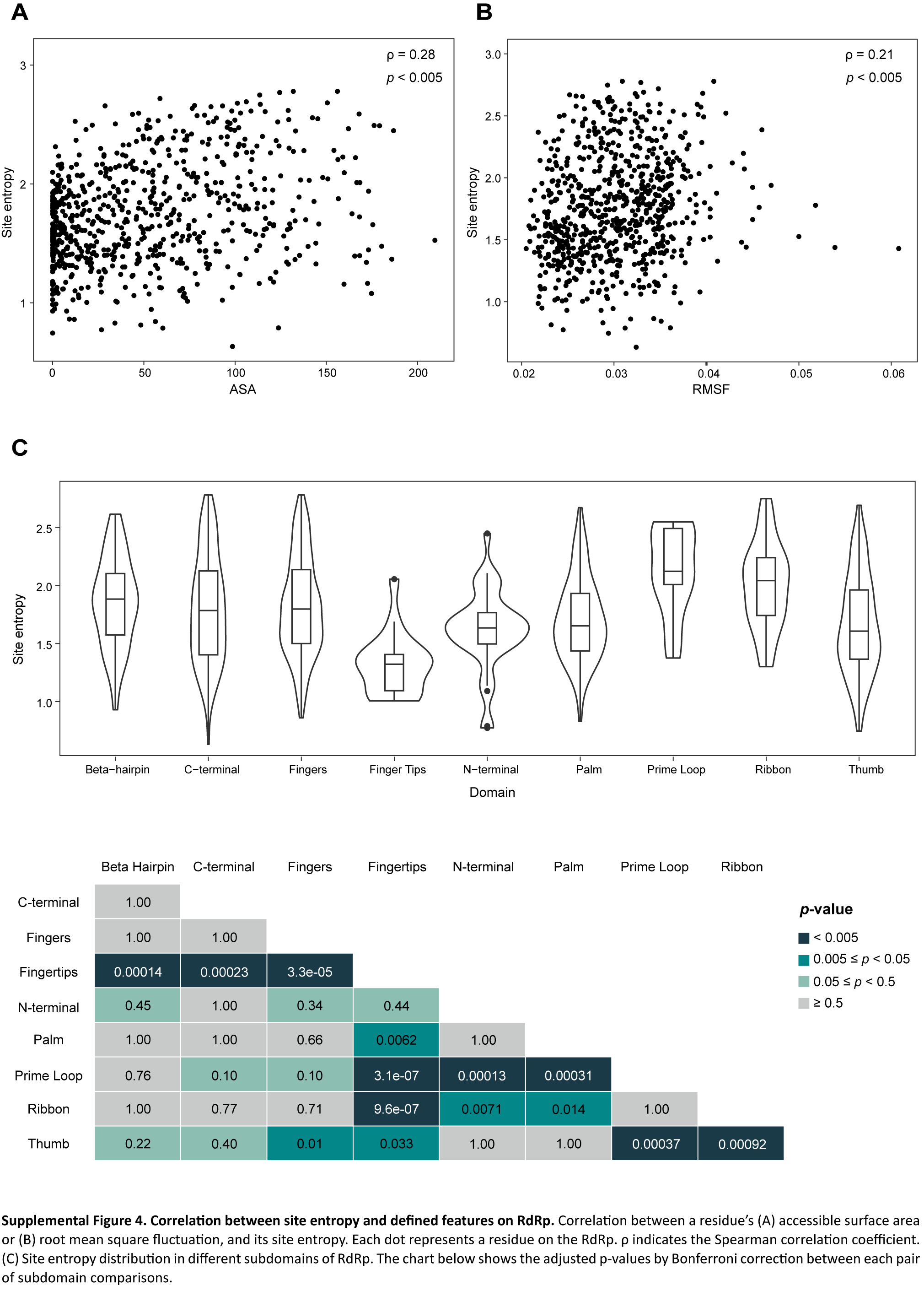

### Supplemental Figure 5

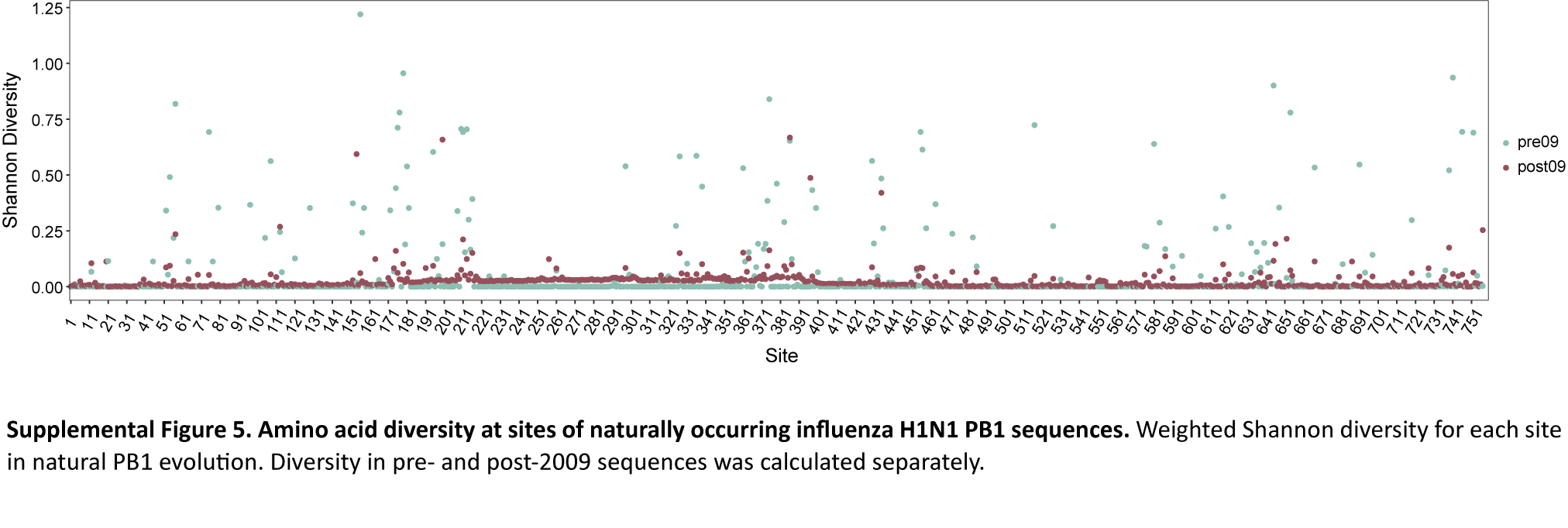

### Supplemental Figure 6

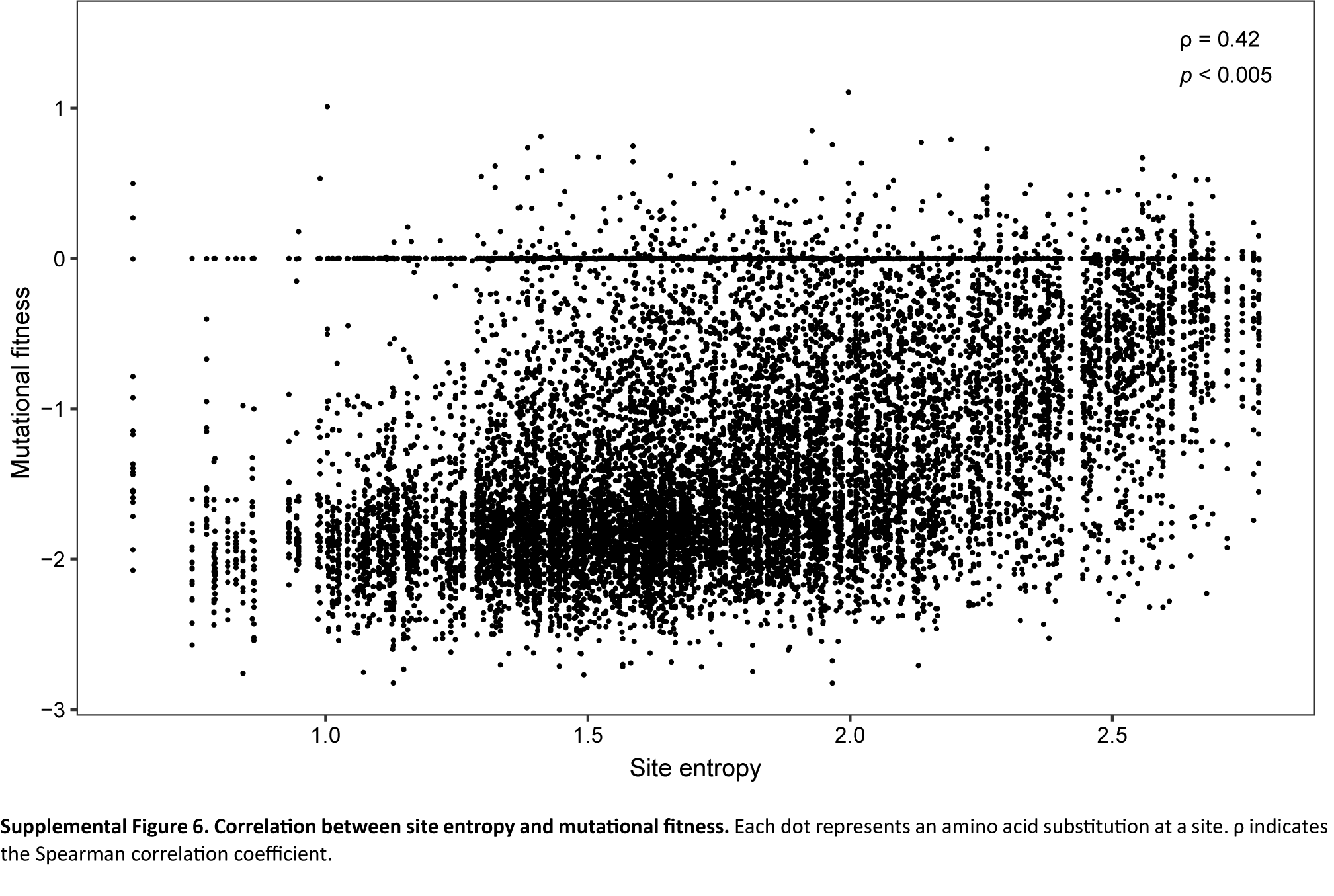

### Supplemental Figure 7

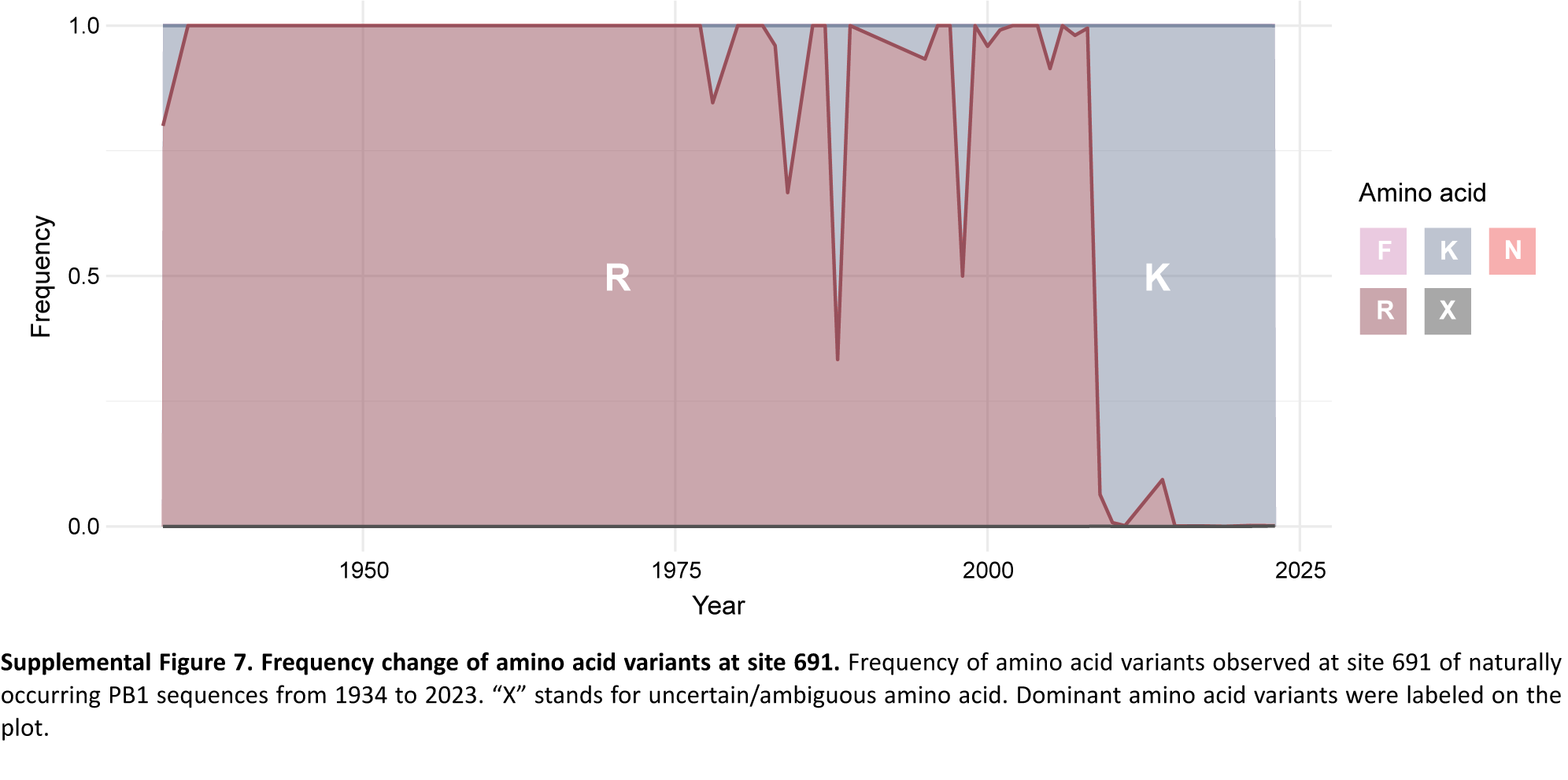
