## Supplemental Text for "Deep mutational scanning reveals the functional constraints and evolutionary potential of the influenza A virus PB1 protein"

**Primers**

**PCR0**

| Rnd0_Fwd | AGCGAAAGCAGGCAAACCATTTGA |
| --- | --- |
| Rnd0_Rev | GGCATTTTTTCATGAAGGACAAGCTAAATTCA |

**PCR1**

| Rnd1_Fwd1 | CTTTCCCTACACGACGCTCTTCCGATCTNNNNNNNNAGCGAAAGCAGGCAAACCATTTGA |
| --- | --- |
| Rnd1_Rev1 | GGAGTTCAGACGTGTGCTCTTCCGATCTNNNNNNNNACCAGGATGGGATTCCTCAAGGAA |
| Rnd1_Fwd2 | CTTTCCCTACACGACGCTCTTCCGATCTNNNNNNNNGATTGTGTATTGGAAGCAATGGCC |
| Rnd1_Rev2 | GGAGTTCAGACGTGTGCTCTTCCGATCTNNNNNNNNCTTAGTCATATTGTCTCTCACTCG |
| Rnd1_Fwd3 | CTTTCCCTACACGACGCTCTTCCGATCTNNNNNNNNGAGATCACAACTCATTTTCAGAGAAAGAGA |
| Rnd1_Rev3 | GGAGTTCAGACGTGTGCTCTTCCGATCTNNNNNNNNGAAATTTCAGTGTCCTGAGAATTGGT |
| Rnd1_Fwd4 | CTTTCCCTACACGACGCTCTTCCGATCTNNNNNNNNGGCAAATGTTGTAAGGAAGATGATG |
| Rnd1_Rev4 | GGAGTTCAGACGTGTGCTCTTCCGATCTNNNNNNNNCATTCCAGGGCTCAATGATGC |
| Rnd1_Fwd5 | CTTTCCCTACACGACGCTCTTCCGATCTNNNNNNNNCGGCCGCTCTTAATAGATGGGACT |
| Rnd1_Rev5 | GGAGTTCAGACGTGTGCTCTTCCGATCTNNNNNNNNGCTGGGAAGCTCCATGCTGAAATT |
| Rnd1_Fwd6 | CTTTCCCTACACGACGCTCTTCCGATCTNNNNNNNNTTCTATCGTTATGGGTTTGTTGCC |
| Rnd1_Rev6 | GGAGTTCAGACGTGTGCTCTTCCGATCTNNNNNNNNGTATAAATTTGGGCCTCCGTC |
| Rnd1_Fwd7 | CTTTCCCTACACGACGCTCTTCCGATCTNNNNNNNNAAAGCTGGACTGCTGGTCTCC |
| Rnd1_Rev7 | GGAGTTCAGACGTGTGCTCTTCCGATCTNNNNNNNNGTTGCAGCACTTTTGGTACATTTG |
| Rnd1_Fwd8 | CTTTCCCTACACGACGCTCTTCCGATCTNNNNNNNNGCCAAAGAGGAATACTTGAAGATGAA |
| Rnd1_Rev8 | GGAGTTCAGACGTGTGCTCTTCCGATCTNNNNNNNNGGCATTTTTTCATGAAGGACAAGCTAAATTCA |

**PCR2**

| Rnd2_Fwd1 | AATGATACGGCGACCACCGAGATCTACACTCGTGGAGCGACACTCTTTCCCTACACGACGCTCTTCCGATCT |
| --- | --- |
| Rnd2_Rev1 | CAAGCAGAAGACGGCATACGAGATCGCTCAGTTCGTGACTGGAGTTCAGACGTGTGCTCTTCCGATCT |
| Rnd2_Fwd2 | AATGATACGGCGACCACCGAGATCTACACCTACAAGATAACACTCTTTCCCTACACGACGCTCTTCCGATCT |
| Rnd2_Rev2 | CAAGCAGAAGACGGCATACGAGATTATCTGACCTGTGACTGGAGTTCAGACGTGTGCTCTTCCGATCT |
| Rnd2_Fwd3 | AATGATACGGCGACCACCGAGATCTACACTATAGTAGCTACACTCTTTCCCTACACGACGCTCTTCCGATCT |
| Rnd2_Rev3 | CAAGCAGAAGACGGCATACGAGATATATGAGACGGTGACTGGAGTTCAGACGTGTGCTCTTCCGATCT |
| Rnd2_Fwd4 | AATGATACGGCGACCACCGAGATCTACACACCAGCGACAACACTCTTTCCCTACACGACGCTCTTCCGATCT |
| Rnd2_Rev4 | CAAGCAGAAGACGGCATACGAGATTCGTCTGACTGTGACTGGAGTTCAGACGTGTGCTCTTCCGATCT |
| Rnd2_Fwd5 | AATGATACGGCGACCACCGAGATCTACACCATACACTGTACACTCTTTCCCTACACGACGCTCTTCCGATCT |
| Rnd2_Rev5 | CAAGCAGAAGACGGCATACGAGATGAACATACGGGTGACTGGAGTTCAGACGTGTGCTCTTCCGATCT |
| Rnd2_Fwd6 | AATGATACGGCGACCACCGAGATCTACACTCGGCAGCAAACACTCTTTCCCTACACGACGCTCTTCCGATCT |
| Rnd2_Rev6 | CAAGCAGAAGACGGCATACGAGATAACCATTCTCGTGACTGGAGTTCAGACGTGTGCTCTTCCGATCT |
| Rnd2_Fwd7 | AATGATACGGCGACCACCGAGATCTACACCTAATGATGGACACTCTTTCCCTACACGACGCTCTTCCGATCT |
| Rnd2_Rev7 | CAAGCAGAAGACGGCATACGAGATGGTTGCCTCTGTGACTGGAGTTCAGACGTGTGCTCTTCCGATCT |
| Rnd2_Fwd8 | AATGATACGGCGACCACCGAGATCTACACGGTTGCCTCTACACTCTTTCCCTACACGACGCTCTTCCGATCT |
| Rnd2_Rev8 | CAAGCAGAAGACGGCATACGAGATCTAATGATGGGTGACTGGAGTTCAGACGTGTGCTCTTCCGATCT |
| Rnd2_Fwd9 | AATGATACGGCGACCACCGAGATCTACACCGCACATGGCACACTCTTTCCCTACACGACGCTCTTCCGATCT |
| Rnd2_Rev9 | CAAGCAGAAGACGGCATACGAGATTCGGCCTATCGTGACTGGAGTTCAGACGTGTGCTCTTCCGATCT |
| Rnd2_Fwd10 | AATGATACGGCGACCACCGAGATCTACACGGCGAGATGGACACTCTTTCCCTACACGACGCTCTTCCGATCT |
| Rnd2_Rev10 | CAAGCAGAAGACGGCATACGAGATTTCTATGGTTGTGACTGGAGTTCAGACGTGTGCTCTTCCGATCT |
| Rnd2_Fwd11 | AATGATACGGCGACCACCGAGATCTACACAATAGAGCAAACACTCTTTCCCTACACGACGCTCTTCCGATCT |
| Rnd2_Rev11 | CAAGCAGAAGACGGCATACGAGATCCTCGCAACCGTGACTGGAGTTCAGACGTGTGCTCTTCCGATCT |
| Rnd2_Fwd12 | AATGATACGGCGACCACCGAGATCTACACTCGTATGCGGACACTCTTTCCCTACACGACGCTCTTCCGATCT |
| Rnd2_Rev12 | CAAGCAGAAGACGGCATACGAGATATGTCGTGGTGTGACTGGAGTTCAGACGTGTGCTCTTCCGATCT |
| Rnd2_Fwd13 | AATGATACGGCGACCACCGAGATCTACACGTCGATTACAACACTCTTTCCCTACACGACGCTCTTCCGATCT |
| Rnd2_Rev13 | CAAGCAGAAGACGGCATACGAGATCGTATAATCAGTGACTGGAGTTCAGACGTGTGCTCTTCCGATCT |
| Rnd2_Fwd14 | AATGATACGGCGACCACCGAGATCTACACAGTGGTCAGGACACTCTTTCCCTACACGACGCTCTTCCGATCT |
| Rnd2_Rev14 | CAAGCAGAAGACGGCATACGAGATGCCATTAGACGTGACTGGAGTTCAGACGTGTGCTCTTCCGATCT |

**Cycling Programs**

**PCR0**

| **Step** | **Temperature(℃)** | **Time(sec)** |
| --- | --- | --- |
| 1. polymerase activation | 95 | 120 |
| 2. denaturing | 95 | 20 |
|  | 70 | 1 |
| 3. annealing | 52 | 30 |
| 4. extension | 70 | 50 |
| Repeat step 2-4 | For 22 cycles in total | |
| 5. hold | 4 | |

**PCR1**

| **Step** | **Temperature(℃)** | **Time(sec)** |
| --- | --- | --- |
| 1. polymerase activation | 95 | 120 |
| 2. denaturing | 95 | 20 |
|  | 70 | 1 |
| 3. annealing | 54 | 20 |
| 4. extension | 70 | 20 |
| Repeat step 2-4 | For 9 cycles in total | |
| 5. termination | 95 | 60 |
| 6. hold | 4 | |

**PCR2**

| **Step** | **Temperature(℃)** | **Time(sec)** |
| --- | --- | --- |
| 1. polymerase activation | 95 | 120 |
| 2. denaturing | 95 | 20 |
|  | 70 | 1 |
| 3. annealing | 55 | 20 |
| 4. extension | 70 | 20 |
| Repeat step 2-4 | For 24 cycles in total | |
| 5. hold | 4 | |
